# Distinct responses of oomycete plant parasites according to their lifestyle in a landscape-scale metabarcoding survey

**DOI:** 10.1101/2020.04.05.026096

**Authors:** Anna Maria Fiore-Donno, Michael Bonkowski

**Affiliations:** Terrestrial Ecology Group, Institute of Zoology, University of Cologne, Cologne, Germany; (AMFD), (MB); Cluster of Excellence on Plant Sciences (CEPLAS), Cologne

**Keywords:** Oomycota, biogeography, soil protists, environmental filtering, plant pathogens, functional traits

## Abstract

Oomycetes are an ubiquitous protistan lineage including devastating crop parasites. Although their ecology in agrosystems has been widely studied, little is known of their distribution in natural and semi-natural ecosystems. We provide here a baseline of the diversity and distribution of soil oomycetes, classified by lifestyles (biotrophy, hemibiotrophy and saprotrophy), at the landscape scale in temperate grassland and forest. From 600 soil samples, we obtained 1,148 Operational Taxonomy Units representing ∼20 million Illumina reads (region V4, 18S rRNA gene). We found a majority of hemibiotrophic plant pathogens, which are parasites spending part of their life cycle as saprotrophs after the death of the host. Overall both grassland and forest constitute an important reservoir of plant pathogens. In forests, relative abundances of obligate biotrophs and hemibiotrophs differed between regions and showed opposite responses to edaphic conditions and human-induced management intensification, suggesting different ecological requirements for these two functional guilds.

## 1. Introduction

### 1.1 General introduction to oomycetes: Taxonomic and functional composition and ecological requirements

Oomycetes are ubiquitous and widespread in terrestrial (Singer et al. 2016; Lara and Belbahri 2011; Geisen et al. 2015), freshwater (Duffy et al. 2015) and marine ecosystems (Garvetto et al. 2018). As protists, they are included in the superphylum Stramenopiles (or Heterokonta) in the superkingdom Harosa (or “SAR”) (Ruggiero et al. 2015). They include c. 2,000 known species, mostly in the two crown groups, the Saprolegniales and Peronosporales (95% of the species) (Thines and Choi 2016). In terrestrial ecosystems, oomycetes occur as pathogens (obligate biotrophs or hemibiotrophs) on plants and other eukaryotes and, less commonly, as saprotrophs (Marano et al. 2016), the plant pathogens representing more than 60% of the oomycete taxa (Thines and Kamoun 2010). Well-known examples are the soil-borne downy mildews with genera like *Phytophthora* and *Pythium* and the white rusts (*Albugo*) on plant leaves (Savory et al. 2015). The genus *Pythium* is one of the most important soil-borne plant pathogens, being ubiquitous and with an extremely wide host range, attacking the roots of thousands of different plant species (Beakes and Thines 2016). *Phytophthora*, the “plant destroyer”, is responsible for the widespread rapid tree decline (Hayden et al. 2013) and for damages to important crops like soybean, tomato, grapevine and tobacco (Lebeda et al. 2008). Because of their negative economic impact, oomycetes are well-studied in silico and as many as 67 species have their genome available in genomes in public databases (https://www.ncbi.nlm.nih.gov/genome, last accessed 28 March 2020). Although they affect forest ecosystems worldwide (Packer and Clay 2000) and are also common pathogens in grasslands (Foley and Deacon 1985), their occurrence in natural habitats, despite of their ecological role in maintaining plant species diversity (Bever et al. 2015) is still poorly explored compared to agrosystems.

### 1.2 Description of the lifestyles

The main distinction among oomycete lifestyles lies between saprotrophs, biotrophs and hemibiotrophs. Free-living saprotrophs live on organic matter or dead tissues. Lifestyles of pathogens include obligate biotrophy which requires living plant tissue (e.g. *Lagena, Peronospora* and *Plasmopara*) and hemibiotrophy (e.g. *Phytophthora* and *Pythium*) characterized by an initial biotrophic infection, followed by necrotrophy on killed host tissue (Pandaranayaka et al. 2019; Ah-Fong et al. 2019). It is important to note that these categories describe life phases rather than the microorganisms themselves (Lorang 2019). Many hemibiotrophic intermediate states between biotrophy and necrotrophy are recognized (Spanu and Panstruga 2017), varying in the length of the latent period, the degree of inflicted damage and the susceptibility of the host (Précigout et al. 2020). In addition, most species of hemibiotrophs can live as saprotrophs in the soil, even in the absence of the host plant (Lifshitz and Hancock 1983), and .recently, species of *Pythium* have been shown to play an important role in litter degradation (Kramer et al. 2016). It is now widely recognized that the ancestral lifestyle was saprotrophy (Spanu and Panstruga 2017), followed by transitions to necrotrophy and hemibiotrophy/biotrophy (Fletcher et al. 2018), so that the traits necessary to overcome the immune system of the host were the last to be acquired. Biotrophs originated repeatedly from hemibiotrophic lineages (Thines and Kamoun 2010; Fletcher et al. 2018). Their complex life cycle, including a stage with large, mycelium-like structures protected by cell walls, the occurrence of motile cells for dispersion, and thick-walled resting stages (Beakes and Thines 2016), may explain their adaptability and dispersal potential.

### 1.3 What is know about the distribution of oomycetes

Recent large-scale studies, using next-generation sequencing have revealed that Oomycota represented c. 5-10% of the protistan metatranscriptomics in forest and grassland soils and in a lesser extent in peatlands (Geisen et al. 2015). A study conducted only on peat bogs, using ad-hoc primers, revealed nevertheless the occurrence of 34 phylotypes, most of which could not be assigned to known species (Singer et al. 2016). Few plant-associated oomycetes were found in the roots of oaks (Sapp et al. 2019), in that of *Arabidopsis thaliana* (Sapp et al. 2018) and in various other plant roots along a glacier chronosequence (Dickie et al. 2019). However, the few dozen of phylotypes found in roots are in contrast with the number of recorded species, suggesting that a wider diversity should be detectable by metabarcoding of oomycetes in soil, where they can persist as saprophytes or as resting spores (Vetukuri et al. 2020).

### 1.4 Introducing our study

Our aim was to assess the biodiversity of oomycetes in a large-scale environmental survey in grasslands and forests in the Biodiversity Exploratories in Germany, and the influence of edaphic, environmental and anthropogenic factors on the oomycete communities, classified according to their lifestyle (obligate biotrophs, hemibiotrophs or saprotrophs) and substrate preference (animal- or plant-associated). Providing a detailed baseline data on oomycete taxonomic and trophic modes distribution is an important contribution to the understanding of ecological processes and ecosystem functioning.

## 2. Results

### 2.1 Sequencing

We obtained more than 22 million reads per run. During the sequencing the overall quality was good, between 84 and 92% ≥ Q30. The reads had on average a length of c. 240 bp, and the overlap during assemblage was of c. 200 bp. In total, we obtained 1,148 oomycete OTUs representing 20,501,201 sequences (Table 1) in 600 grasslands and forest sites, with on average 34,168 reads/site (minimum 460, maximum 108,910). Our primers also sequenced c. 28% of non-oomycete taxa (mostly sequences that could not be identified in the database). Because of the high variability of the ITS 1 region, no satisfactory automated alignment could be obtained, and the chimera check had to be performed with non-aligned sequences, a somewhat less stringent method, probably leaving a proportion of undetected chimeras in our database, although c. 7% were deleted (Table 1). The 1,148 OTUs represented 319 unique Blast best hits. Only 39% of the OTUs were 96-100% similar to any known sequence (Fig. S1). The 30 most abundant OTUs (> 10,000 sequences) accounted for 73% of the total sequences, while 526 OTUs of < 1,000 sequences contributed only to 1,6 % of all sequences (Table S2). A database with the OTU abundance per sample, taxonomic assignment and estimated functional traits is provided (Table S2 and Table S3). Our sequencing and sampling efforts were sufficient, since the actual richness was reached after only 310,000 sequences (Fig. S2A) and at 200 samples (Fig. S2B), so that the observed distribution patterns would not have been influenced by undersampling.

**Table 1.** Quality estimates and error rate for each run; initial, assembled, quality-trimmed, and unique number of reads for each run. Number of reads retrieved at each step of the sequence processing for the 4 assembled runs.

| Illumina runs | >Q30<br>% | %<br>error<br>rate | reads | assembled<br>reads | % assembled<br>reads | trimmed | %<br>unique<br>reads | 4 runs<br>assembled,<br>total reads | clusters<br>(=OTUs) | OTUs -<br>rare | representing<br>sequences | OTUs<br>genuine | OTUs non-<br>chimeric | representing<br>sequences |
| --- | --- | --- | --- | --- | --- | --- | --- | --- | --- | --- | --- | --- | --- | --- |
| Grassland 2011 | 84 | 2.7 | 24'342'940 | 11'987'411 | 49 | 4'816'104 | 28 | 23'920'438 | 350'045 | 1'717 | 22'473'536 | 1'238 | 1'148 | 20'501'201 |
| Grassland 2017 | 92 | 1.55 | 22'687'371 | 11'958'891 | 53 | 6'516'916 | 34 |  |  |  |  |  |  |  |
| Forest 2011 | 87 | 1.7 | 25'228'346 | 12'202'548 | 48 | 6'507'303 | 26 |  |  |  |  |  |  |  |
| Forest 2017 | 88 | 1.6 | 28'070'312 | 13'579'536 | 48 | 6'080'115 | 37 |  |  |  |  |  |  |  |

### 2.2 Diversity of oomycetes

At a high taxonomic level, the majority of the OTUs could be assigned to the Peronosporales (73%) and only 21% to the Saprolegniales, with Leptomitales (5%) and Haptoglossales (1%) only marginally present, the latter probably because our primers did not match all of them (Fig. 1). We did not use the family rank because the traditional classification is not supported by most phylogenies and varies between authors. Among the 32 identified genera, the most common was by far *Pythium* (50%), followed by *Saprolegnia* (10%), *Aphanomyces* (6%), *Phytophthora* (5%) and *Apodachlya* (5%). The Haptoglossales (1%) contain only a dozen species, all of which are parasites of rotifers and bacterivorous nematodes (Beakes and Thines 2016). Representative of the order Rhipidiales (largely saprotrophic and aquatic) are missing from the reference database (Supplementary information S1) and were thus not being found in our samples.

**Fig. 1.**
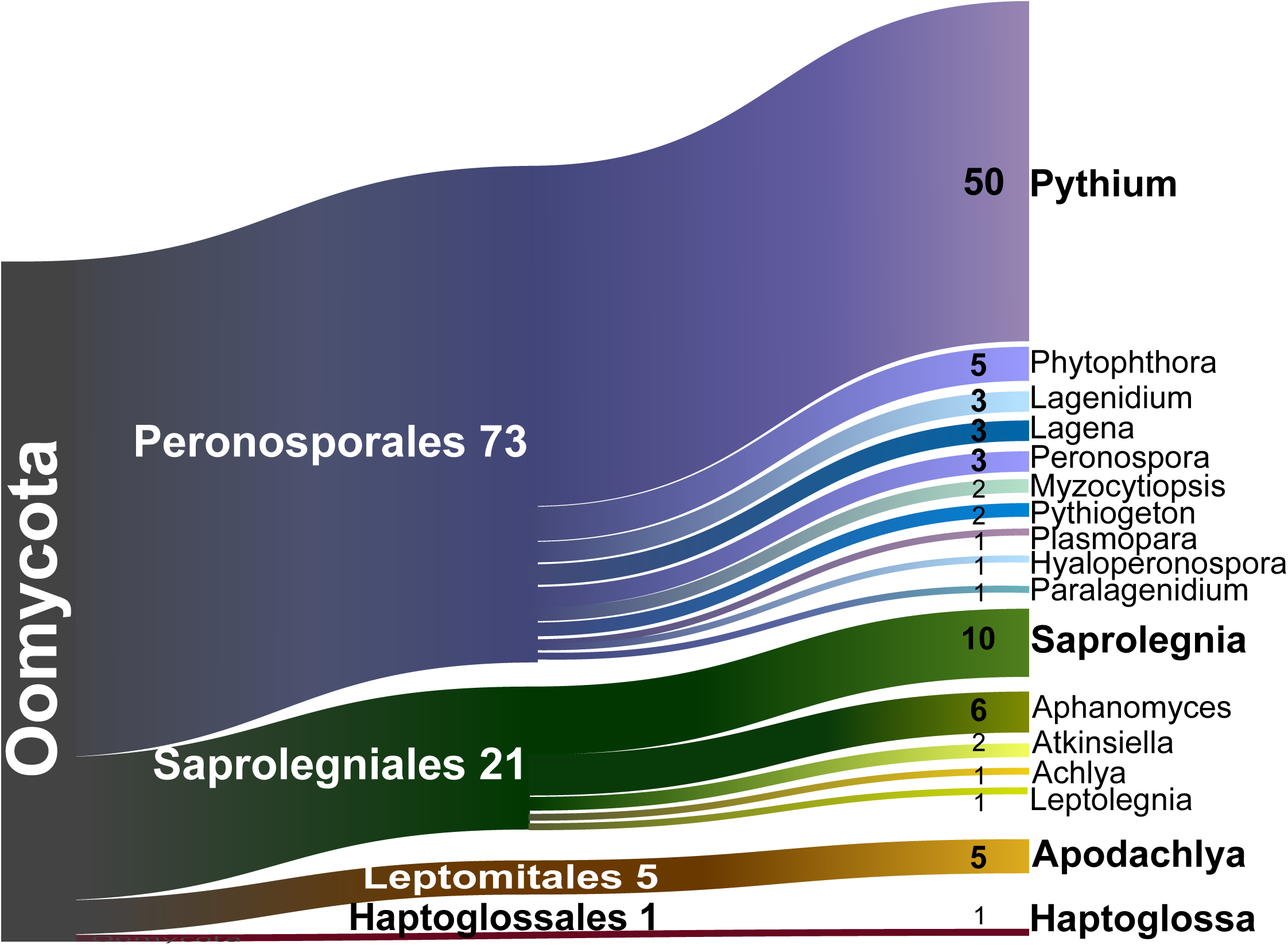
Sankey diagram showing the relative contribution of the OTUs to the taxonomic diversity. Taxonomical assignment is based on the best hit by BLAST. From left to right, names refer to phyla, orders (ending -ida) and genera. Numbers are percentages of OTUs abundance - OTUs representing <1% are not shown

### 2.3 Functional diversity

The majority of the OTUs were hemibiotrophs (75-76%, 860 OTUs), that is, spending part of their lifetime as parasites of the living host and part as saprotrophs after the death of the host. The most abundant genera were pathogens of plants, i.e. *Pythium* (67% of the hemibiotrophs) and *Phytophthora* (6%). Abundant genera of animal pathogens were *Saprolegnia* (14%), *Lagenidium* (5%), *Myzocytiopsis* (3%), *Atkinsiella* (2%) (Fig. 1, Fig. 2 & Table S2). The second category, far below in percentage (10-11%) were the obligate biotrophs, mostly of plants, e.g. *Lagena* (34% of the obligate biotrophs), *Peronospora* (28%), *Plasmopara* (14%) and *Hyaloperonospora* (9%). The saprotrophs, e.g. *Apodachlya* (70% of the saprotrophs) and *Leptolegnia* (14%), counted only for 6-8% of all OTUs. The relative proportions of each lifestyle were quite similar between forest and grassland (Fig. 2). Most oomycetes depended on plants (65%), more in grassland (69%) than in forest (66%). Animal-dependent oomycetes (21% in total) were slightly more abundant in forest (21%) than in grassland (17%) (Fig. 2).

**Fig. 2.**
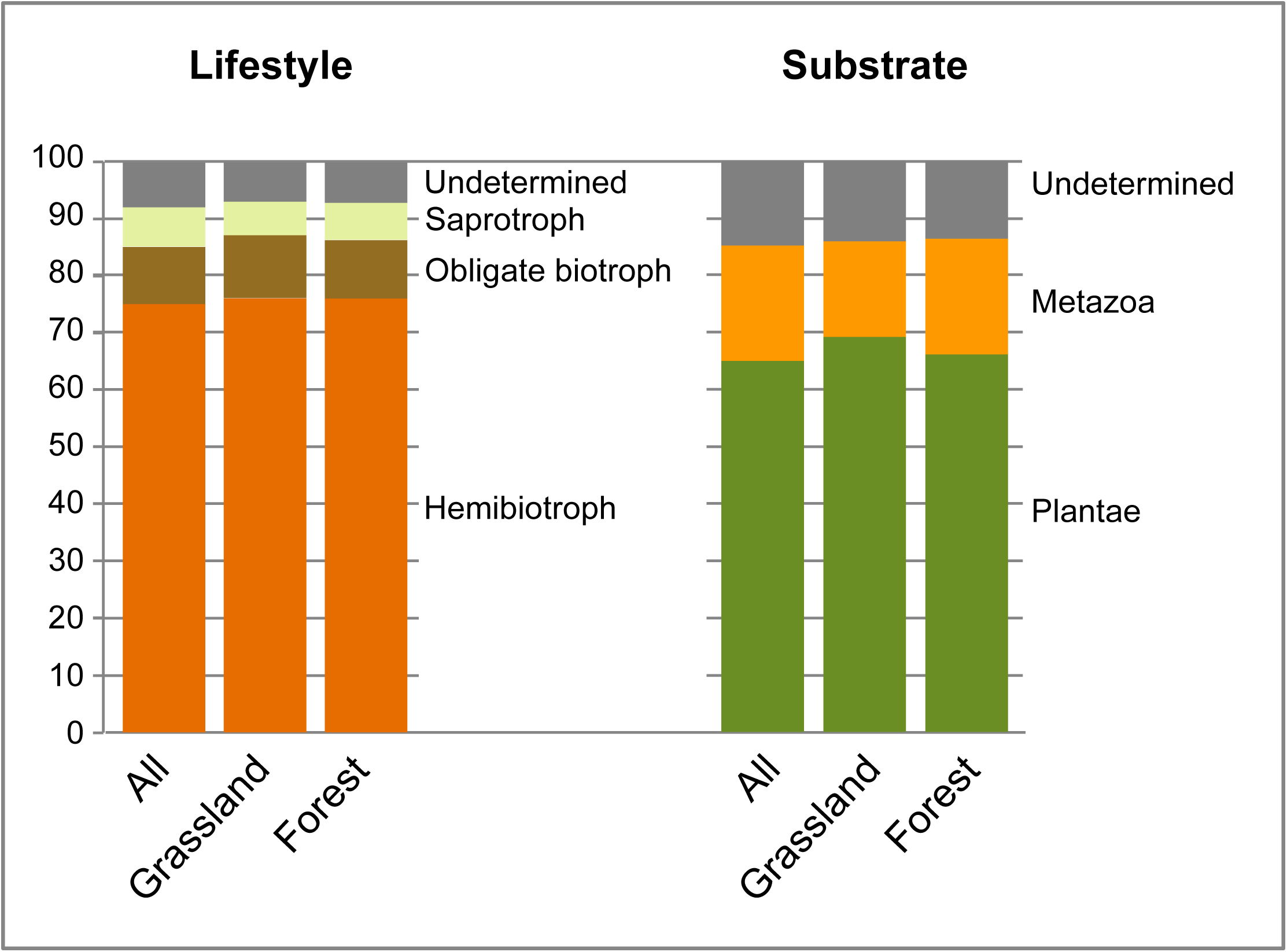
Histograms showing the relative contribution of the OTUs to the functional diversity. Total number of OTUs: all=1,148; grassland=1,007; forest=1,072. Lifestyle and substrate are determined according to Tables S2-S3. OTUs representing <1% are not shown

### 2.4 Alpha and beta diversity

A total of 1,007 and 1,072 oomycete OTUs were retrieved in grassland and forest sites, respectively. 931 OTUs (81%) were shared between ecosystems, with 76 unique to grassland and 141 to forest. Most OTUs were shared between regions, with only 37 missing in Schorfheide, five in Hainich and eight in Alb (Table S2). Alpha diversity, as revealed by Shannon indices, was significantly higher in grassland than in forest, while at the regional level, only Schorfheide had a significantly lower alpha diversity. Evenness of oomycete OTUs was higher in grassland than in forest, and higher in Alb and Hainich than in Schorfheide (Fig. S3). Despite the almost ubiquitous presence of oomycete OTUs in grasslands, PCoA revealed differences between communities. Both PCoA components, explaining 34% and 13% of the variance in Bray-Curtis distances, showed gradients from grassland to forest, although with some overlap (Fig. 3). The grassland communities were more similar between themselves than the forest communities, where the effect of the region was more pronounced, with Schorfheide standing out, especially its forest communities (Fig. 3).

**Fig. 3.**
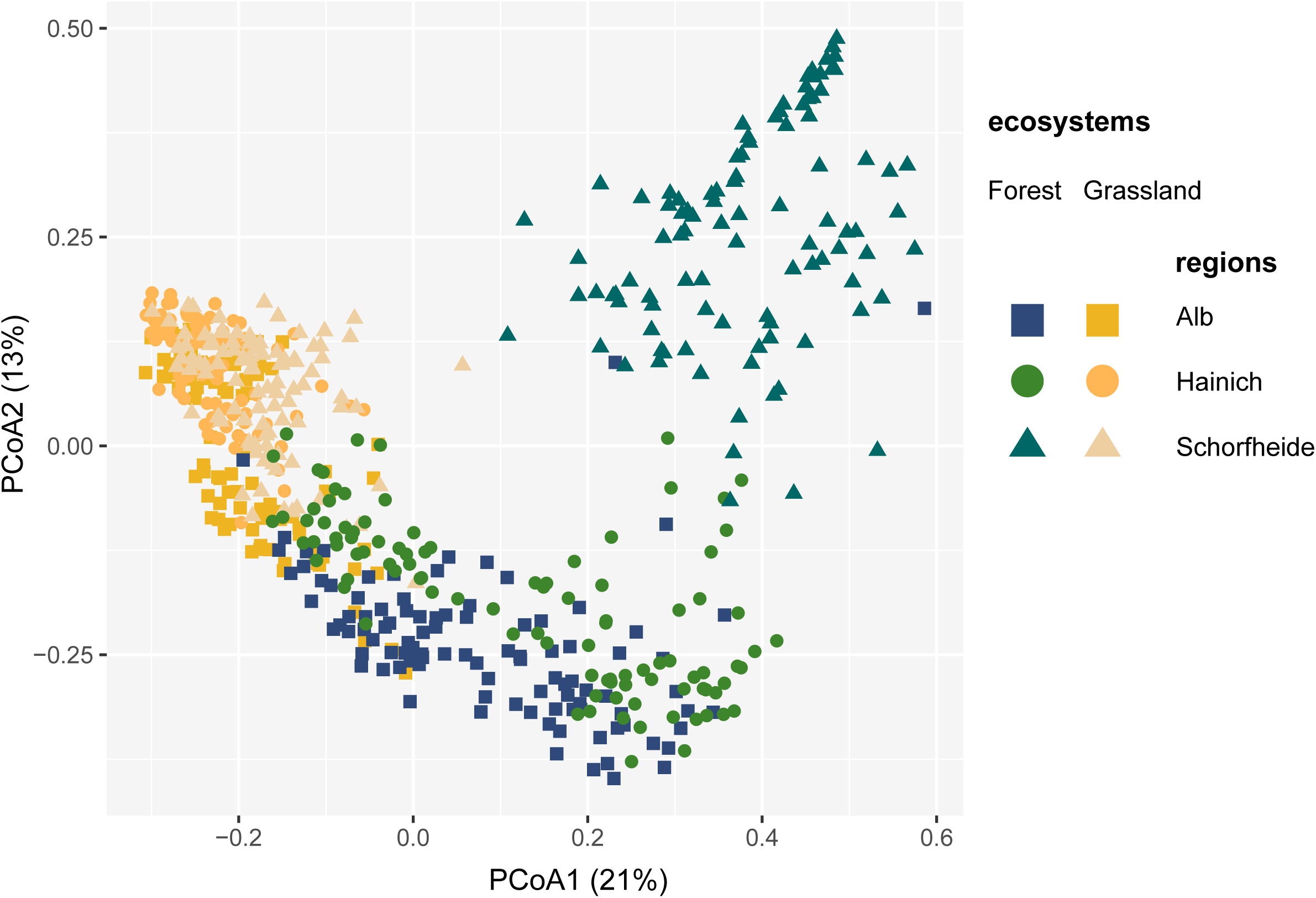
Principal Component Analysis of the Bray-Curtis dissimilarity indices between oomycete OTUs, showing grassland sites clustering together, while forest sites are more region-driven, especially Schorfheide

**Fig. 4.**
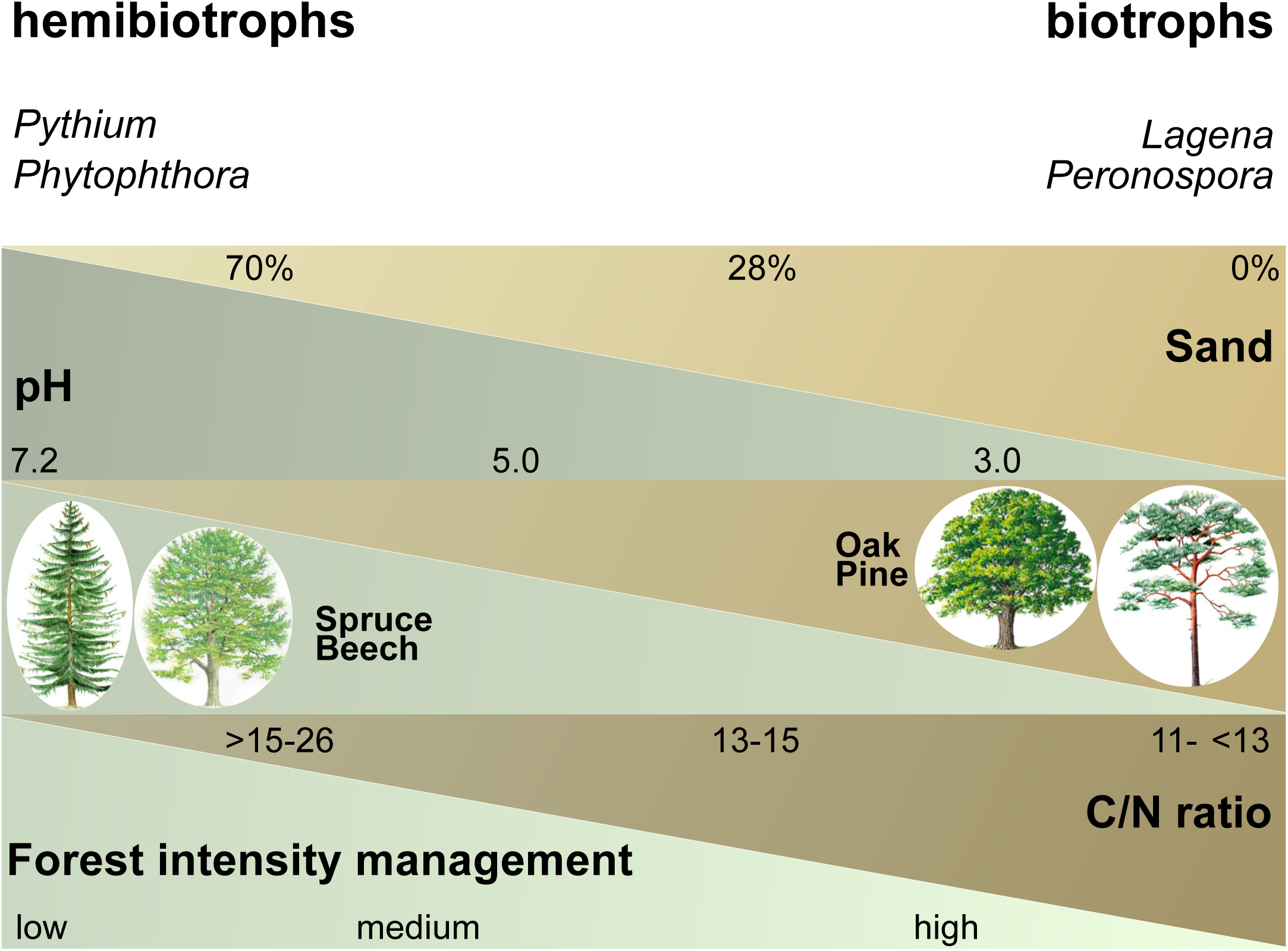
Schematic illustration showing the positive or negative influences of the most influential ecological and edaphic parameters in forests (selected by the models in Table S5) and the relative abundances of the oomycetes lifestyles. For numeric parameters, minimum, average and maximum values are given. Forest developmental stage was estimated by the mean of the diameter of 100 trunks of well-developed trees of the dominant species

### 2.5 Major drivers of species turnover in oomycete communities

Variation partitioning among the three predictors indicated that ecosystem (grassland vs forest), region (Alb, Hainich and Schorfheide) and year of sampling (2011 and 2017) together accounted for 28.6% (adjusted R^2^) of the total variation in oomycete beta diversity. Ecosystem explained 16.3% of the variation, region 10.0%, and the year of sampling only 2.2%. Among the soil parameters that were measured in the framework of the Biodiversity Exploratories (Table S4 & Table S5), carbon content (total, organic and inorganic) and nitrogen (total N) were all co-correlated; they were tested separately, and based on a slightly higher significance in the models, organic carbon was kept. Among the three co-correlated soil components, sand was preferred over clay or silt, because of its recognized importance for oomycetes occurrence. The soil and ecological parameters included in our analyses could explain more variance in forests (R^2^ 10.5-25.3) than in grasslands (R^2^ 2.6-5.9) (Table S6). The most parsimonious models identified an influence of soil type in grassland as well as in forest (Table S6). Other explanatory factors in grassland, but only for specific lifestyles or regions, were the LUI index, mowing intensity, the soil C/N ratio and the organic C content. In addition to soil type, more significant explanatory factors were identified in forest: main tree species, soil pH, sand content, forest management intensity. Other factors, selected by a minority of the models, were forest developmental stage, organic C content, and C/N ratio (Table S6)..

We further investigated the effects of these selected environmental parameters on each of the oomycete lifestyles, by comparing their relative abundances in each ecosystem. Hemibiotrophs and biotrophs (and to a lesser extent, also saprotrophs) showed opposite responses to environmental parameters, both in grassland (Fig. S4) and forest (Fig. S5). Less significant differences were found in grassland than in forest, since the OTUs in the former communities were more evenly distributed (Fig. 3). The oomycete communities in grasslands of Schorfheide stood out in respect to Hainich, with slightly decreasing biotrophs and increasing saprotrophs (Fig. S4A), a tendency also partially reflected by the communities of specific soil types that were either only present in Hainich or in Schorfheide (Fig. S4B). Hemibiotrophs decreased at a higher land use intensity (LUI index) (Fig. S4C). An increasing C/N ratio, indicative of soils poor in nitrogen, led to a relative decrease of hemibiotrophs and an increase of biotrophs (Fig. S4D). In soils with high organic carbon content, biotrophs decreased and saprotrophs increased (Fig. S4E).

In forest, hemibiotrophs and biotrophs together with saprotrophs showed opposite shifts in relative abundances patterns. At the regional scale, Schorfheide differed from Hainich and Alb by a decrease of hemibiotrophs and an increase of biotrophs and saprotrophs (Fig. S5A), a trend mirrored by the soil types unique to each region (Fig. S5B) and further by the tree species unique to Schorfheide, i.e. pine and oak (Fig. S5C). Hemibiotrophs were reduced, while biotrophs and saprotrophs slightly increased with increasing forest management intensity and increasing sand content (Fig. S5D & E). With an increasing C/N ratio, hemibiotrophs decreased and biotrophs increased (as in grasslands) but also saprotrophs increased (Fig. S5F). Hemibiotrophs decreased in more acidic soils (mostly present in Schorfheide), while biotrophs and saprotrophs increased (Fig. S5G).

## 3. Discussion

Our data stemmed from a thorough sampling across three regions spanning a South-North gradient in Germany using taxon-specific primers. Based on the high sequencing depth (saturation was reached) (Fig. S2) we could detect detailed responses of each lifestyle to the ecological processes involved in shaping their distribution. The high percentage of OTUs not closely matching any published sequence (Fig. S1) suggests a significant hidden species richness not yet taxonomically recorded or sequenced in the ITS1 database. Consistently with previous studies on protists, a high alpha-diversity and a low beta-diversity were found (Fiore-Donno et al. 2019; Lentendu et al. 2018). Our data not only confirm the low endemicity of oomycetes, but thank to our thorough sampling, high sequencing depth and the use of taxon-specific primers, we showed that almost all OTUs were shared between ecosystems and regions. This implies that community assembly of oomycetes is not limited by dispersal over countrywide distances. As a corollary, the remarkable differences in beta diversity are probably explained by contrasted responses of lifestyles to environmental selection (Fiore-Donno et al., 2019).

### 3.1 Dominance of plant-associated hemibiotrophs in natural and semi-natural ecosystems; significance for agriculture and ecosystem dynamics

We found that plant-associated hemibiotrophic oomycetes dominated in natural and semi-natural ecosystems, which thus constitute a reservoir of plant pathogens with the potential to spread to neighbouring agrosystems. *Pythium* and *Phytophthora*, the two most infamous destructive oomycete plant pathogens, together constituted 73% of the hemibiotrophs in grassland and forest. Half of the OTUs and 67% of the hemibiotrophs were attributed to the genus *Pythium* (Fig. 1), a result in line with previous oomycete community studies (Sapkota and Nicolaisen 2015; Riit et al. 2016; Venter et al. 2017), and by Peronosporales being by far the largest order in Oomycota, comprising more than 1,000 species (Thines 2014). The ability to alternate between saprotrophy and parasitism likely favoured the hemibiotrophs over the obligate biotrophs, which in our study represented only 10% of the OTUs (Fig. 2). The obligate biotrophic downy mildews were represented by *Peronospora* (3%, c. 400 described species), which parasitizes a wide range of flowering plants, the Brassicaceae-infecting *Hyaloperonospora* (1%) and *Plasmopara* with a wide host spectrum in eudicots (Thines and Choi 2016). *Lagena* (3%), a root-infecting parasite of grasses, including cereals (Blackwell 2011) was also the most frequent OTU in plant-root associated oomycetes (Dickie et al. 2019).

The oomycete communities were more evenly distributed in grassland than in forest. Grassland soil have a higher microbial biomass than forests (Dequiedt et al. 2011). In our study, differences between grassland and forest, in terms of OTUs richness, alpha diversity and evenness, were small. In a study comparing ecotypes across Wales, the relative abundance of oomycetes compared to other Stramenopiles (algal Ochrophyta, mostly) decreased from crops and grasslands to forests and bogs - the absolute numbers of oomycete OTUs were not provided (George et al. 2019). This is in contrast with reports on Oomycetes representing the dominant Stramenopiles in forest soils (Geisen et al. 2015). Although we could not distinguish between soil- and air-borne species, the leaf-infecting white blister rusts (mainly the genus *Albugo*), which are obligate angiosperm parasites (Beakes and Thines 2016), were nearly absent from your soil study, with only two OTUs attributed to *Pustula*. Their absence in 600 soil samples confirmed that they mostly spread by air or possibly by vertical transmission as seed endophytes (Ploch and Thines 2011). Our study, in assessing the biogeography of oomycete pathogens at a national scale, also contributes to a better understanding of the mechanisms of dispersal from natural reservoirs to agrosystems, and perhaps could contribute to better reveal the evolutionary processes leading to the emergence of new pathogens.

### 3.2 Obligate biotrophs and hemibiotrophs showed opposite responses in forests

In forests, hemibiotrophs and obligate biotrophs showed opposite responses to a number of environmental factors, suggesting different ecological requirements of both functional groups. In summary, relative abundance of hemibiotrophs was decreased in nitrogen-poor, sandy and acidic soils, which are the type of soils common in Schorfheide, planted with pines and oaks (Fig. S5). The same combination of edaphic and ecological factors which led to a reduced abundance of hemibiotrophs favoured biotrophs and saprotrophs. This was a surprising result: we expected hemibiotrophs to show intermediate responses, while obligate parasites and saprotrophs would each represent opposite extremes of the response gradient. This assumption was based on the evolutionary distance - saprotrophy being the ancestral stage and biotrophy the last to evolve - and the substrate preferences. Our data clearly show that environmental factors act differentially on hemibiotrophs versus obligate biotrophs and saprotrophs.

Soil texture is well known to influence the abundance and disease development of oomycetes. Sandy soils are less prone than clayey soils to suffer temporal waterlogging conditions that strongly benefit the pathogens (Gómez-Aparicio et al. 2012), so that an increasing proportion of sand in soil had a negative effect on pathogen abundance (Gómez-Aparicio et al. 2012). Accordingly, we found that hemibiotrophs strongly decreased in high sand content soils in forest (Fig. S5E), while soils periodically inundated (Stagnosols) had high relative abundances (Fig. S5B). Soils with a high organic content, or soils amended with compost, have long been known to suppress a number of *Pythium* and *Phytophthora* species (Hayden et al. 2013), but the reason is not precisely known. One hypothesis is that carbon-rich soils harbour a wide diversity of microbes that may outcompete oomycetes, especially hemibiotrophic species with saprotrophic capacities, like some *Pythium* (Hayden et al. 2013). Not only carbon, but also nitrogen content has an effect on soil suppressiveness, which occurs less frequently under high-nutrient conditions (Löbmann et al. 2016). Accordingly, relative abundances of hemibiotrophs, both in grassland and forest, decreased in nitrogen-poor soils with a high carbon content (but not that of biotrophs) (Figs. S4D & S5F), while saprotrophs, being carbon-dependent, increased with C/N ratio in forest (Fig. S5F) and with organic C content in grassland (Fig. S4E). Thus, in order to successfully enhance soil suppressiveness, it is necessary to understand how particular management practices will differentially influence each key component of biodiversity (Löbmann et al. 2016; Schlatter et al. 2017). Here, we show that a more intensive forest management, as recorded in the Biodiversity Exploratories (including estimation of harvested trunks, invasive tree species and cut dead wood) will not increase the abundance of the hemibiotrophs.

### 3.3 Saprotrophs and aquatic taxa in soils

More puzzling was the presence in soil of “water-moulds” (Saprolegniales, 21%), although their constant and widespread occurrence in soils in addition to aquatic habitats has been documented in the past (Dick and Newby 1961; Johnson et al. 2002). Some species of *Saprolegnia* are known as fish or fish eggs parasites (saprolegniosis in fisheries), but are also saprotrophs (Johnson et al. 2002). Some species of *Atkinsiella* are known for the damages they inflict on marine crustaceans, but there are also parasites of terrestrial invertebrates (Beakes and Thines 2016). The supposed predominance of the genus *Aphanomyces* (Beakes and Thines 2016) is not fully confirmed by our study (6% of the OTUs). Species of the genus can be saprotrophs or hemibiotrophs, so in our classification it is reported as “undetermined lifestyle”.

### 3.4 Other factors influencing oomycete communities

The distribution of oomycete communities is only partially explained in our study by ecosystem, region or year of collection (only 29% of the variation explained). We hypothesized that plant diversity and evenness and the presence of suitable plant hosts (especially for oomycetes with a narrow host range) may be explanatory factors that were not recorded: vegetation is a major determinant of the spatial distribution of soil pathogens (Gómez-Aparicio et al. 2012). Microbial ecology studies suffer from a lack of systematic basic information on community distribution which hampers predicting effects of anthropogenic environmental changes. Providing a detailed baseline data on the occurrence of oomycete taxa, the ecology and the distribution of their lifestyles is an important contribution to the understanding of ecological processes and ecosystem functioning, a prerequisite for subsequent analyses linking them to the distribution and diversity of potential plant hosts.

## 4. Materials and Methods

### 4.1 Study site, soil sampling and DNA extraction

Our study took place in three German Biodiversity Exploratories, i.e. the Biosphere Reserve Schorfheide-Chorin in the State of Brandenburg, the National Park Hainich and its surroundings in the State of Thuringia and the Biosphere Reserve Schwäbische Alb in the State of Baden-Württemberg (Fischer et al. 2010). Each exploratory comprises 50 grassland sites from extensive pastures to highly fertilized meadows and 50 differently managed forest sites, mainly beech (*Fagus sylvatica*) sometimes mixed with spruce (*Picea abies*). Each site contains a study plot of 20 × 20 m. From all study plots, 300 soil samples were collected in a coordinated joint sampling campaign within 14 days in April 2011 and a second one in April 2017. From each plot 14 soil cores of 8.3 cm diameter were taken every 3 m along two transects of 20 m each, oriented North-South and East-West, employing a soil corer. The surface layer (0-10 cm) was collected, after removing plants, pebbles and conspicuous roots. Soil cores from each plot were sieved (2 mm mesh size), mixed, homogenised and immediately frozen for further analysis. Soil DNA was extracted from 400 mg of soil, 3- to 6-times, using the DNeasy PowerSoil Kit (Qiagen GmbH, Hilden, Germany) following the manufacturer’s protocol, to obtain a sufficient amount to be shared between research groups of the Biodiversity Exploratories.

### 4.2 and barcodes design, amplification

We designed primers to target the ITS1, because it better allows to distinguish between species (although not in the closely related species of some groups of the polyphyletic *Pythium*) (Steciow et al. 2014), and its reference database is rich (c. 600 species). Potential problems with the ITS1 marker are the variation in length (216-534 bp, the longest sequences in *Plasmopara*). Because the Oomycota are a large and diverse assemblage, we could design primers to match the two major clades (in number of species and ecological importance), the Saprolegniales and the Peronosporales, missing the small basal lineages, i.e. the marine Eurychasmales, Haliphthorales and Olpidiopsidales and partially the nematode infecting Haptoglossales. We discarded other potential markers such as the V4 region of the small subunit of the ribosomal RNA gene because it does not display enough intraspecific variation, and the mitochondrial cytochrome oxidase cox2 because it has a very high AT rich content, which could negatively interfere with the quality of the Illumina run.

Primers were designed using an alignment of 2,941 Oomycetes ITS1 sequences. To build it, we downloaded all ITS sequences >500 bp from GenBank RefSeq (accessed 24.08.2017), clustered them at 96% similarity using cd-hit 454 (Beifang et al. 2010). This database has been used for Blast purposes (Supplementary information S1). For primer design, we removed sequences with ambiguities, aligned them with MAFFT (Katoh and Standley 2013); the alignment was refined by hand using Bioedit (Hall 1999) and cut to the fragment of interest. Our forward primers were overlapping the position of several primers at the end of the SSU, e.g. ITS1-ofw (Liebe et al. 2016), ITS1oo (Riit et al. 2016) and OOMUP18Sc (Lievens et al. 2004); the reverse primer in the 5.8S gene was partially overlapping primers ITS1-orw (Liebe et al. 2016) and ITS-3oo (Riit et al. 2016). We designed three primers (two forward primers partially overlapping) to be used in two successive semi-nested PCRs. The primers did not form any hairpin, primer-dimers with themselves or with each other, and their hybridization temperatures were similar. The length of the targeted fragment varied from 206 to 534 bp. The first PCR was conducted with the primers S1777F (non-specific - 5’ GGTGAACCTGCGGAAGGA 3’) (1777 is the starting position in the SSU sequence *Saccharomyces cerevisiae* Z75578) and 58SOomR (oomycete-specific - 5’ TCTTCATCGDTGTGCGAGC 3’). The second PCR was conducted with barcoded primers S1786StraF (specific for Stramenopiles - 5’ GCGGAAGGATCATTACCAC 3’) - and the 58SOomR as before. The barcodes consisted in eight-nucleotide-long sequences appended to the 5’-ends of both the forward and the reverse primers, because tagging only one primer leads to extensive mistagging (Esling et al. 2015). To design the barcodes, we first used barcrawl (Frank 2009) to obtain a list of barcodes with a balanced nucleotide content (no homopolymers), not folding on themselves and to themselves an the attached primer (no “hairpin”), not forming heteroduplexes with the corresponding primer and having at least 3 bases differences between them. In addition, using custom R scripts, we selected from the previous list the barcodes that did not match the consensus of the reference alignment flanking the primer region (forward: GTGAACCT, reverse: VCGCTGCG) and without cross-dimerization between each combination of primer+barcodes. We designed 18 barcoded versions for the forward and the reverse primers allowing for 324 possible combinations to label samples of which only 150 were used, since it is advisable to leave a proportion of unused combinations to decrease mistagging (Esling et al. 2015). Barcoded primers were specifically ordered for NGS application to Microsynth (Wolfurt, Austria) (Table S1).

For the amplification, we incorporated 1 µl of 1:10 soil DNA template for the first PCR and 1 µl of the resulting amplicon as a template for a following semi-nested PCR. We employed the following final concentrations: GreenTaq polymerase (Fermentas, Canada) 0.01units, buffer 1x, dNTPs 0.2 mM and primers 1 µM. The thermal programme consisted of an initial denaturation step at 95°C for 2 min, 24 cycles at 95°C for 30 s, 52°C for 30 s, 72°C for 30 s; and a final elongation step at 72°C for 5 min. The number of PCR cycles was kept at 24 since chimera formation arises dramatically after 25 cycles (Michu et al. 2010). All PCRs were conducted twice to reduce the possible artificial dominance of few amplicons by PCR competition (2 × 10 µl for the first and 2 × 27 µl for the second PCR), and the two amplicons were pooled after the second PCR.

### 4.3 Library preparation and sequencing

The amplicons were checked by electrophoresis and 25 μl of each were purified and normalized using SequalPrep Normalization Plate Kit (Invitrogen GmbH, Karlsruhe, Germany) to obtain a concentration of 1-2 ng/μl per sample, which were then pooled, totalling four samples (forest 2011 and 2017 and grassland 2011 and 2017). During the library preparation amplicons were end-repaired, small fragments were removed, 3’ ends were adenylated, and Illumina adapters and sequencing primers were ligated (TruSeqDNA PCR-Free, Illumina Inc., San Diego, CA, USA). The library was quantified by qPCR, performed following the manufacturer’s instructions (KAPA SYBR® FAST qPCR Kit, Kapabiosystems, Wilmington, MA, USA) on a CFX96 Real Time System (Bio-Rad, Hercules, CA, USA). Sequencing was performed with a MiSeq v3 Reagent kit of 600 cycles on a MiSeq Desktop Sequencer (Illumina Inc., San Diego, CA, USA) at the University of Geneva (Switzerland), department of Genetics and Evolution. Four runs were conducted in total.

### 4.4 Sequences processing

In each run, paired reads were assembled using mothur v.3.7 (Schloss et al. 2009) (which was also used in the following steps) allowing no difference in the primers and barcodes, no ambiguities and removing assembled sequences < 200 bp, with an overlap <100 bp or with any mismatch in the overlap (Table 1). The quality check and removal/cutting of low-quality reads was conducted with the default parameters. Reads were sorted into samples via detection of the barcodes (Table S1), renamed and then the four runs were assembled. Sequences were clustered using vsearch v.1 (Rognes et al. 2016) in mothur, with abundance-based greedy clustering (agc) and a similarity threshold of 97% (Table 1). Rare OTUs (< 0.001% of the total sequences, in this case <368 reads) were deleted, since we observed that they mostly represented artifactual reads (Fiore-Donno et al. 2018).

Using BLAST+ (Camacho et al. 2008) with an e-value of 1e-^1^ and keeping only the best hit, sequences were identified on a custom ITS database (Supplementary information S1) and non-oomycetes sequences were removed (Table 1). Since the ITS sequences were too variable to be aligned, we performed the chimera check using UCHIME (Edgar *et al*. 2011) as implemented in mothur by taking into account the abundance of the OTUs and the OTUs identified as chimeric were removed. The results are shown as a table with the OTUs abundance/site, their taxonomic assignment according to the best hit by Blast and their functional assignment (Table S2). The relative abundance of each oomycete taxonomic level was illustrated using Sankey diagram generator V1.2 (http://sankey-diagram-generator.acquireprocure.com/, last accessed Jan. 2020) and refined with the open-source vector graphic editor Inkscape (https://inkscape.org/en/, last accessed Jan. 2020).

### 4.5 Functional diversity

We compiled a table with functional traits of ecological importance, i.e. lifestyle and substrate. Oomycetes lifestyles that are traditionally distinguished are saprotrophs - living on dead organic matter (e.g. decaying plants, insect exuviae), obligate biotrophs (parasites or pathogens) and hemibiotrophs (switching from a parasitic to a saprotrophic lifestyle). We also collected information about the substrate - plant or animal - for saprotrophs as well as for pathogens (Table S2). We could only attribute functional traits to the level of the genus, since we clustered the OTUs at 97% similarity, and because the uncertainty of the attribution in the database increases at the species level. We attributed traits by searching in the relevant literature, providing all consulted references (Table S3). We could not find sufficient information about the dispersion mode - aerial, or solely through water and soil.

### 4.6 Statistical analyses

All statistical analyses were carried out within the R environment (R v. 3.5.1) (R Development Core Team 2014), on the OTU abundance/sites table (Table S2), and the environmental parameters (Table S3 & Table S4), the latter normalized by the K-nearest neighbours. Unless otherwise specified, community analyses were performed with the package vegan (Oksanen et al. 2013). *Alpha diversity*: To evaluate if more sampling and sequencing effort would have revealed more richness, we carried out an analysis based on OTUs accumulation curves, function *specaccum*, method rarefaction and 1,000 random permutations; species richness was extrapolated using the function *specpool*. Alpha diversity estimates were based on the relative abundances of OTUs (function *decostand*, method total); Shannon diversity and Pielou’s evenness were obtained with the function *diversity. Beta diversity*: Variation partitioning (function *varpart* applied to the Hellinger-transformed OTUs dataset) was performed to assess the proportion of explained beta diversity by the factors region, year of collection (2011, 2017) or ecosystem (grassland vs forest). Beta diversity between regions and ecosystems was inferred by Principal Coordinate Analysis (PCoA, function *cmdscale*), using Bray-Curtis dissimilarities (function *vegdist*, method “bray”) on the relative abundances of OTUs, then plotted with the package ggplot2. A distance-based redundancy analysis (dbRDA) on Bray-Curtis dissimilarities (function *dbrda*), was used to investigate the effect of environmental factors on the beta-diversity of oomycete communities. Parsimonious models were selected by the function *ordistep* with default parameters based on 999 permutations, and only significant results were shown in dbRDA. All available parameters were tested for co-correlation according to variance inflation factors (function *vif*.*cca*). Among the soil parameters measured in the framework of the Biodiversity Exploratories (Table S4 & Table S5), soil carbon (total, organic and inorganic) and nitrogen (total N) were all co-correlated; based on slightly higher significance in the models organic carbon was kept. Among the three co-correlated descriptors of soil texture, soil sand content was chosen over clay and silt contents.

The effects of environmental parameters on the *relative abundances* of the trophic guilds were tested with general linear models (function *glm*, core package), subjected to the general linear hypothesis test (function *glht*, package multcomp) with Tukey’s test for multiple comparisons of means and a heteroskedasticity-consistent covariance matrix estimation (function *vcovHC*, package sandwich).

## 5. Declarations

### 5.3 Data availability

Raw sequences have been deposited in Sequence Read Archive (NCBI) SRA 10697395-8, Bioproject PRJNA513166, and the 1,148 OTUs (representative sequences) under GenBank accession nos MN268786 - MN269933.

### 5.4 Authors’ contributions

Conceptualization of the PATHOGEN project and interpretation of the data [Bonkowski and Fiore-Donno]. Amplifications, Illumina sequencing, bioinformatics pipeline, statistic analyses and first draft of the manuscript [Fiore-Donno]. Funding acquisition, revisions of the manuscript [Bonkowski].

### 5.5 Ethics approval, Consent to participate, Consent for publication, Code availability

Not applicable.

## 6. Acknowledgments

We are very grateful to Linhui Jiang and Cristopher Kahlich for invaluable help in the lab. We thank Graham Jones, Scotland for writing the R scripts selecting barcodes. At the University of Geneva (CH), we thank Jan Pawlowski and Emanuela Reo and we are liable to the Swiss National Science Foundation Grant 316030 150817 funding the MiSeq instrument. We thank the managers of the Exploratories, Swen Renner, Kirsten Reichel-Jung, Kerstin Wiesner, Katrin Lorenzen, Martin Gorke, Miriam Teuscher, and all former managers for their work in maintaining the plot and project infrastructure; Simone Pfeiffer and Christiane Fischer for giving support through the central office, Jens Nieschulze and Michael Owonibi for managing the central data base, and Markus Fischer, Eduard Linsenmair, Dominik Hessenmüller, Daniel Prati, Ingo Schöning, François Buscot, Ernst-Detlef Schulze, Wolfgang W. Weisser and the late Elisabeth Kalko for their role in setting up the Biodiversity Exploratories project. Field work permits were issued by the responsible state environmental offices of Baden-Württemberg, Thüringen, and Brandenburg.

## Supplementary information

### Supplementary tables

**Table S1**. Combinations of barcodes used in this study, with the corresponding soil samples. Code for sample names: AE= Alb, HE=Hainich, SE=Schorfheide. G=grassland sites - to be replaced by F in forest sites (the same barcodes were applied since the grassland and forest samples were amplified and sequenced separately).

**Table S2**. Database of the abundance of each OTU per sample. The taxonomic assignment (supergroup, class, order, family, genus and species) is provided according to the best hit by BLAST (PR2 database), with the % of similarity given. Functional traits (lifestyle and substrate) were estimated for each genus following Table S3.

**Table S3**. References for the functional traits of the oomycetes genera identified in this study.

**Table S4**. Environmental parameters from the 150 grassland study sites and two years of collection.

**Table S5**. Environmental parameters from the 150 forest study sites and two years of collection.

**Table S6**. Most parsimonious models (RDA), with their respective R^2^ adjusted and F values, and the F values of the elements selected in each model. Significance values shown as symbol (see footnote).

## 9. Supplementary Figures

**Fig. S1** Similarities of the oomycete OTUs with known sequences. OTUs are classified according to their percentage of similarity to the next kin by BLAST. The horizontal bar length is proportional to the number of OTUs in each rank. Shaded area=OTUs with a similarity ≥ 96%

**Fig. S2** Description of the oomycete diversity. **a**. Rarefaction curve describing the observed number of OTUs as a function of the sequencing effort; saturation was reached with c. 310,000 sequences. **b**. Species accumulation curve describing the sampling effort; saturation was reached with 200 samples. In grey, confidence interval

**Fig. S3** Boxplot and table of the alpha diversity of oomycete OTUs estimated with Shannon, inverse Simpson and evenness indices, for all sites and for sites binned by region and ecotype. The highest mean indices are in bold

**Fig. S4** Boxplots of the variation of the oomycete relative abundances of the tree main lifestyles in grassland, coloured according to region. **a**. by region; **b**. by soil type, only Cambisoil is found in the three regions; **c**. by land usage; **d**. by land use intensity (LUI) index, transformed into a categorical variable according to quantiles; **d**. by clay content, transformed as in D. Note that the y-scale varies between graphs. Small red letters: a change from “a” to “b”, or “c” indicates a significant difference (multiple comparison of means, Tukey’s test); two letters (e.g. “ab”) indicate non-significant differences between plots sharing those letters. Red lines indicate the mean

**Fig. S5** Boxplots of the variation of the oomycete relative abundances of the tree main lifestyles in forest, coloured according to region. **a**. by region; **b**. by soil type; **c**. by forest management intensity; **d**. by clay content, transformed into a categorical variable according to quantiles; **e**. by main tree species, Latin names: beech=*Fagus sylvatica*, spruce=*Picea abies*, pine=*Pinus sylvestris*, oak=*Quercus petrea* & *Q. robur*, pine and oak growing only in Schorfheide. **F**. by developmental stage (trunk diameter). Small red letters: a change from “a” to “b”, or “c” indicates a significant difference (multiple comparison of means, Tukey’s test); two or three letters (e.g. “ab” or “abc”) indicate non-significant differences between plots sharing those letters. Red lines indicate the mean

**Supplementary information S1** Reference database of 197,400 ITS sequences used for Blast

