## Supplementary material for "Distinct responses of oomycete plant parasites according to their lifestyle in a landscape-scale metabarcoding survey": Fig. S1

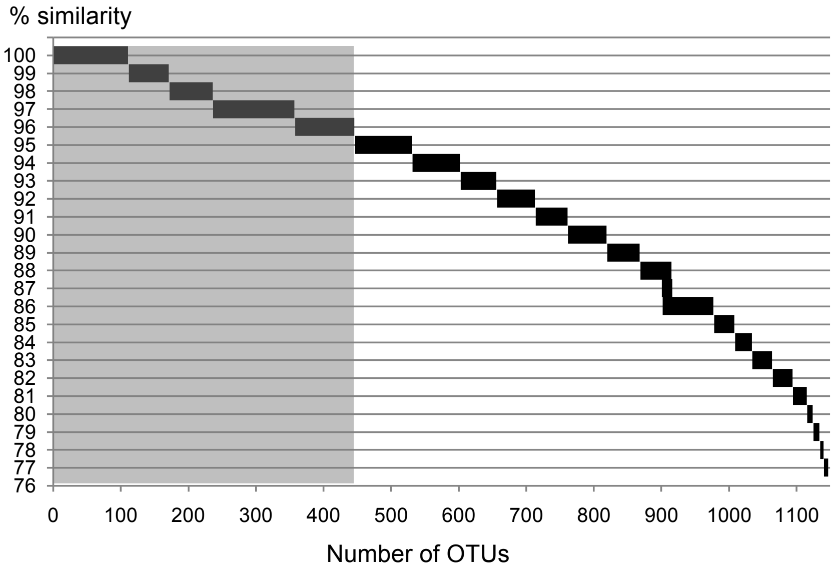


**Fig. S1**. Similarities of the OTUs with known sequences. OTUs are classified according to their percentage of similarity to the next kin by BLAST. The horizontal bar lentgh is proportional to the number of OTUs in each rank. Shaded area=OTUs with a similarity ≥96%
