## Supplementary material for "Distinct responses of oomycete plant parasites according to their lifestyle in a landscape-scale metabarcoding survey": Fig. S2

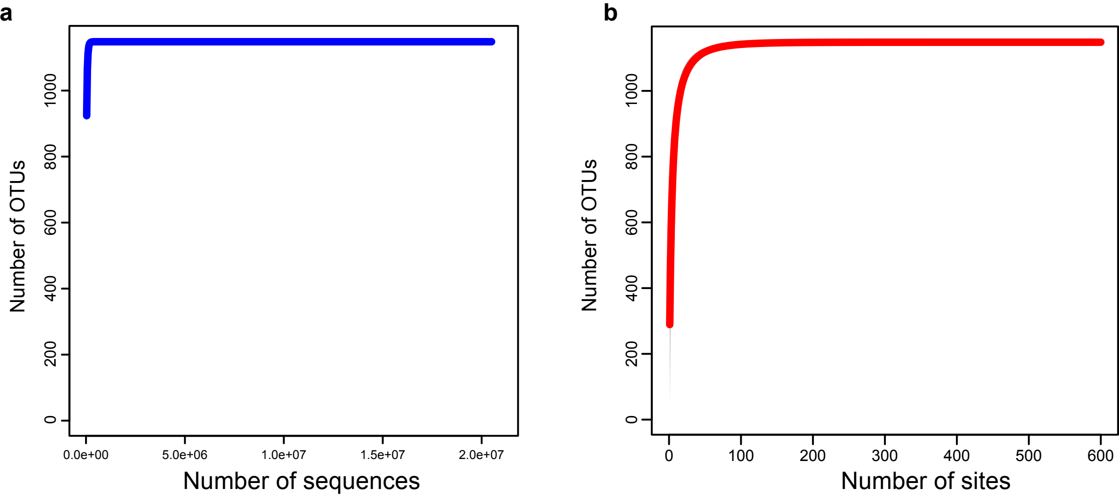


**Fig. S2** Description of the oomycetan diversity. **a**. Rarefaction curve describing the observed number of OTUs as a function of the sequencing effort; saturation was reached with c. 310,000 sequences. **b**. Species accumulation curve describing the sampling effort; saturation was reached with 200 samples. In gray, confidence interval
