## Supplementary material for "Distinct responses of oomycete plant parasites according to their lifestyle in a landscape-scale metabarcoding survey": Fig. S3

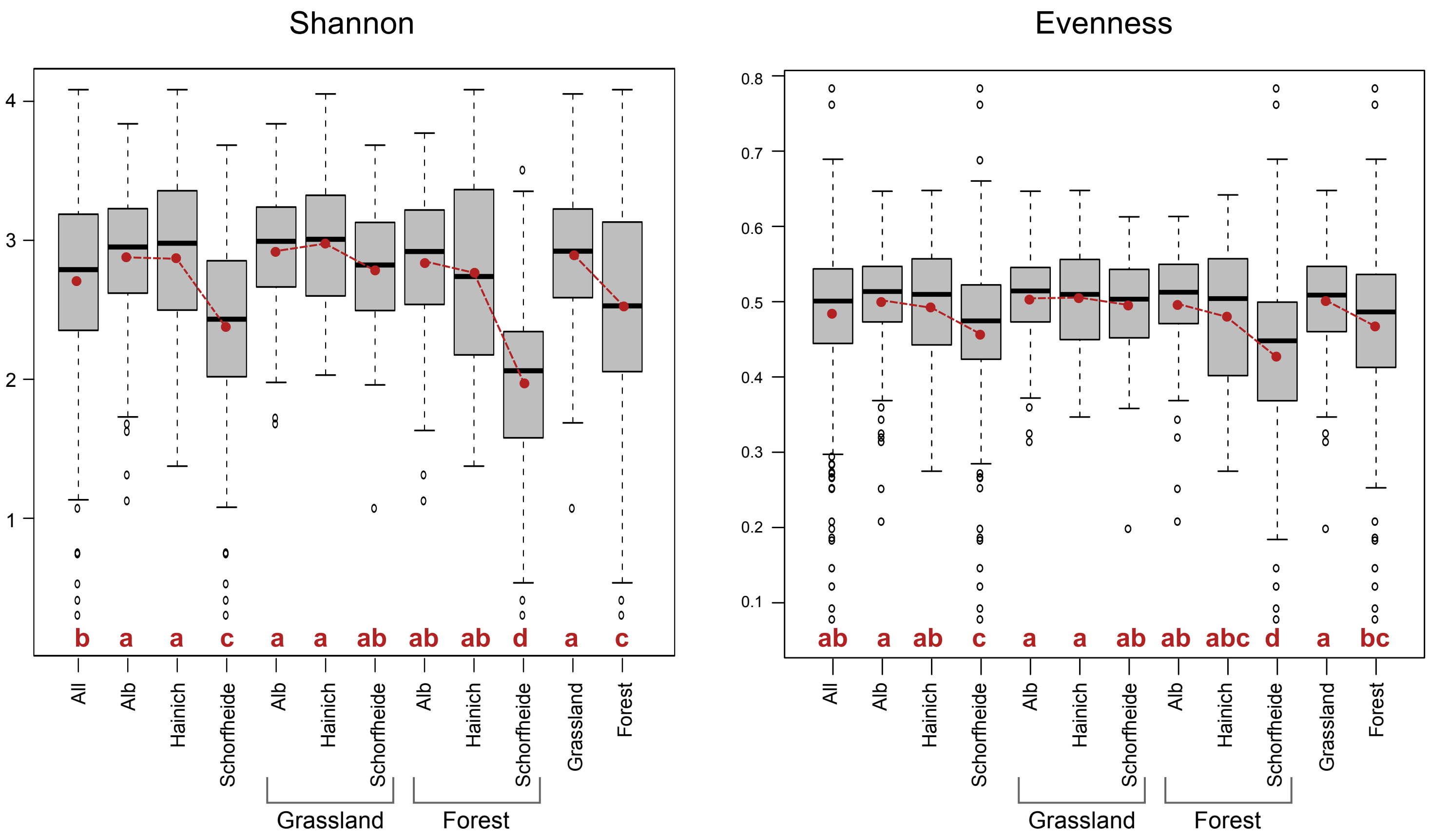


| | All sites | Alb | Hainich | Schorfheide | Alb grassland | Hainich grassland | Schorfheide grassland | Alb forest | Hainich forest | Schorfheide forest | Grassland | Forest |
|---|---|---|---|---|---|---|---|---|---|---|---|---|
| **Shannon** | | | | | | | | | | | | |
| **Min.** | 0.31 | 1.13 | 1.38 | 0.31 | 1.69 | 2.03 | 1.08 | 1.13 | 1.38 | 0.31 | 1.08 | 0.31 |
| **Mean** | 2.72 | **2.89** | **2.89** | 2.39 | 2.93 | **2.99** | 2.8 | **2.85** | 2.78 | 1.98 | 2.91 | 2.54 |
| **Max.** | 4.08 | 3.84 | 4.08 | 3.68 | 3.84 | 4.05 | 3.68 | 3.77 | 4.08 | 3.51 | 4.05 | 4.08 |
| **St_Dev** | 0.62 | 0.48 | 0.58 | 0.66 | 0.45 | 0.44 | 0.43 | 0.51 | 0.67 | 0.59 | 0.45 | 0.71 |
| **evenness** | | | | | | | | | | | | |
| **Min.** | 0.08 | 0.21 | 0.27 | 0.08 | 0.32 | 0.35 | 0.2 | 0.21 | 0.27 | 0.08 | 0.2 | 0.08 |
| **Mean** | 0.49 | **0.5** | 0.49 | 0.46 | **0.51** | **0.51** | 0.5 | **0.5** | 0.48 | 0.43 | **0.5** | 0.47 |
| **Max.** | 0.78 | 0.65 | 0.65 | 0.78 | 0.65 | 0.65 | 0.61 | 0.61 | 0.64 | 0.78 | 0.65 | 0.78 |
| **St_Dev** | 0.09 | 0.07 | 0.08 | 0.1 | 0.06 | 0.07 | 0.06 | 0.07 | 0.09 | 0.12 | 0.06 | 0.1 |

**Fig. S3** Boxplots and table of the alpha diversity of oomycete OTUs estimated with Shannon and evenness indices, for all sites and for sites binned by region and ecotype. Red letters: a change from “a” to “b”, or “c” indicates a significant difference (multiple comparison of means, Tukey's test); two or three letters (e.g. “ab” or "abc") indicate non-significant differences between plots sharing those letters. Red dots indicate the means (black lines the median). In the table, the highest means are in bold
