## Supplementary material for "Distinct responses of oomycete plant parasites according to their lifestyle in a landscape-scale metabarcoding survey": Fig. S4

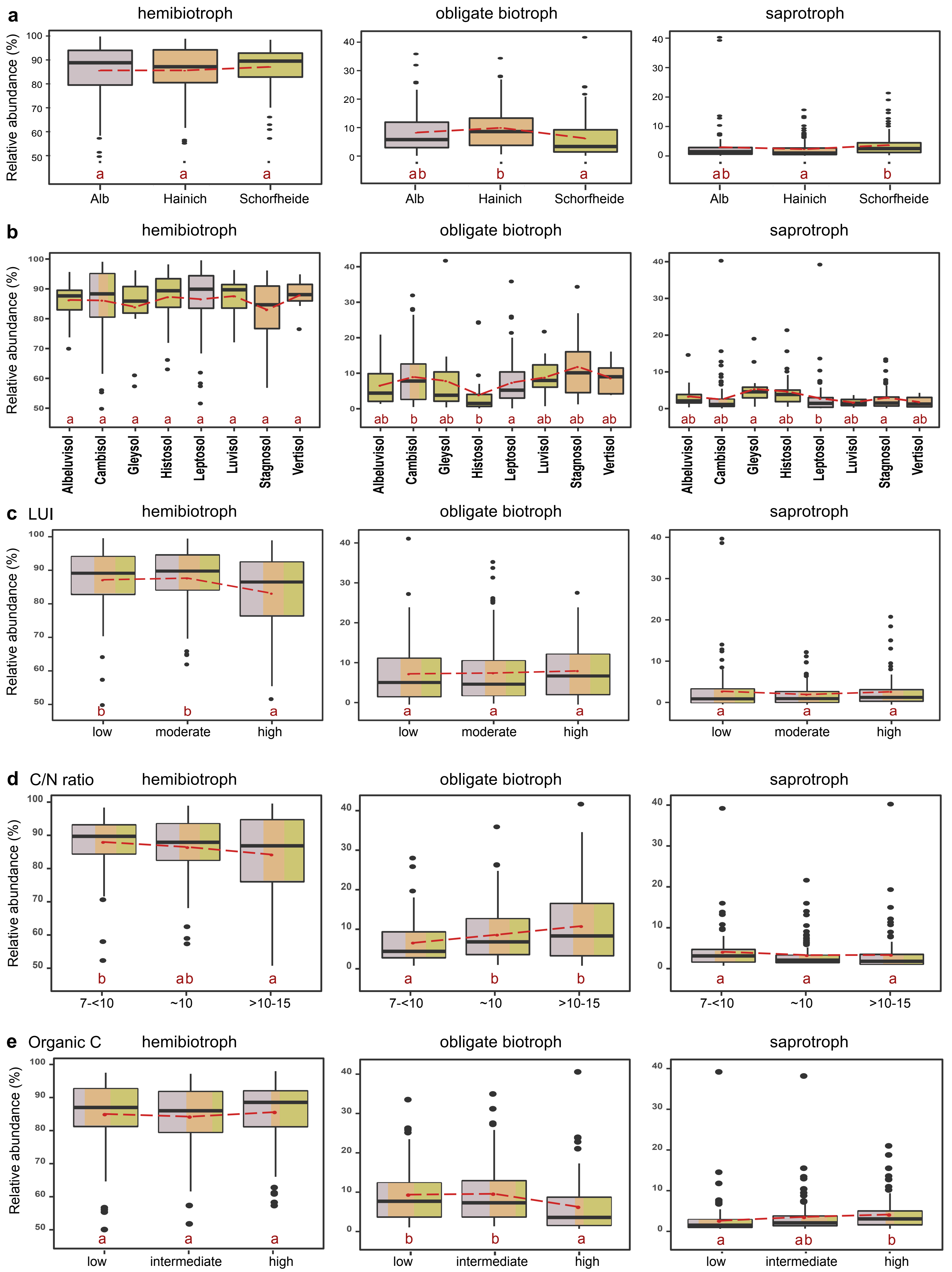


**Fig. S4** Boxplots of the variation of the oomycete relative abundances of the tree main lifestyles in grassland, colored according to region. **a**. by region; **b**. by soil type, only Cambisoil is found in the three regions; **c**. by land use intensity (LUI) index, transformed into a categorical variable according to quantiles; **d**. by C/N ratio, transformed as in c; **e**. by organic carbon content, transformed as in c. The y-scale varies between graphs. Red letters: a change from “a” to “b”, or “c” indicates a significant difference (multiple comparison of means, Tukey's test); two letters (e.g. “ab”) indicate non- significant differences between plots sharing those letters. Red lines indicate the mean
