## Supplementary material for "Distinct responses of oomycete plant parasites according to their lifestyle in a landscape-scale metabarcoding survey": Fig. S5

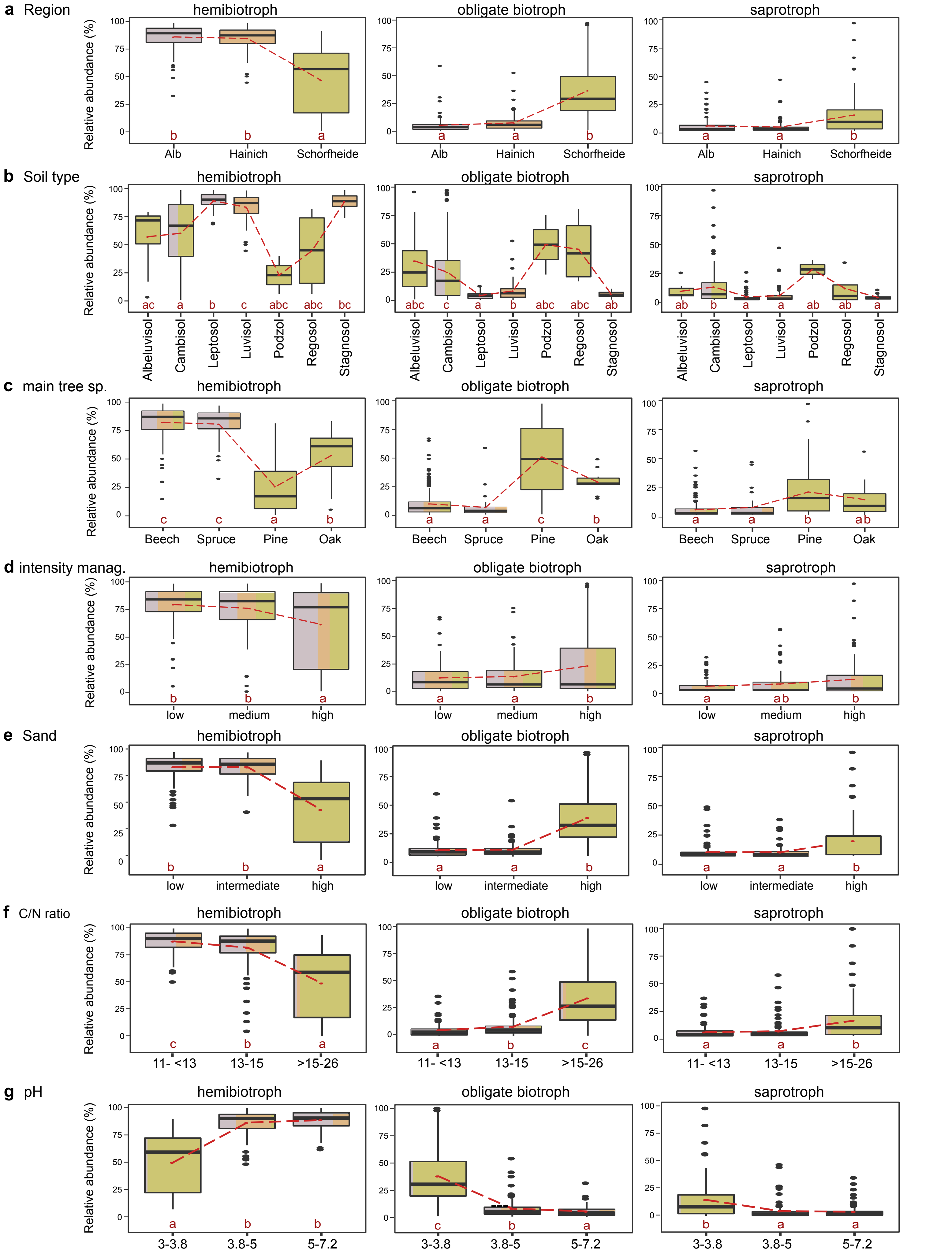


**Fig. S5** Boxplots of the variation of the oomycete relative abundances of the tree main lifestyles in forests, colored according to region. **a**. by region; **b**. by soil type; **c**. by main tree species, Latin names: beech=*Fagus sylvatica*, spruce=*Picea abies*, pine=*Pinus sylvestris*, oak=*Quercus petrea* & *Q. robur,* pine and oak growing only in Schorfheide; **d**. by intensity management; **e**. by sand content, transformed into a categorical variable according to quantiles; **f**. by C/N ratio, transformed as in e; **g**. by pH, transformed as in e. Red letters: a change from “a” to “b”, or “c” indicates a significant difference (multiple comparison of means, Tukey's test); two or three letters (e.g. “ab” or "abc") indicate non-significant differences between plots sharing those letters. Red lines indicate the mean
