## Supplementary material for "Distinct responses of oomycete plant parasites according to their lifestyle in a landscape-scale metabarcoding survey": Table S1

**Table S1**. Combinations of barcodes used in this study, with the corresponding soil samples. Code for sample names: AE=Schwäbische Alb, HE=Hainich, SE=Schorfheide. G stands for grassland sites - to be replaced by W in forest sites (the same barcodes were applied since the grassland and forest samples were amplified and sequenced separatedly).

| Forward | Reverse | Sample | Forward | Reverse | Sample | Forward | Reverse | Sample |
|---|---|---|---|---|---|---|---|---|
| GCTTCTAG | ATAGCTTG | AEG001 | GCTATTGC | CCTTAATG | HEG001 | AGCTAGTA | ATAGCTTG | SEG001 |
| GCTTCTAG | CCTTAATG | AEG002 | GCTATTGC | CACATGCT | HEG002 | AGCTAGTA | CCTTAATG | SEG002 |
| GCTTCTAG | CACATGCT | AEG003 | GCTATTGC | TGTCATGC | HEG003 | AGCTAGTA | CACATGCT | SEG003 |
| GCTTCTAG | TGTCATGC | AEG004 | GCTATTGC | CGATAAGG | HEG004 | AGCTAGTA | TGTCATGC | SEG004 |
| GCTTCTAG | CGATAAGG | AEG005 | GCTATTGC | TTACGCGA | HEG005 | AGCTAGTA | CGATAAGG | SEG005 |
| GCTTCTAG | TTACGCGA | AEG006 | GCTATTGC | TGGAGCTT | HEG006 | AGCTAGTA | TTACGCGA | SEG006 |
| GCTTCTAG | TGGAGCTT | AEG007 | GCTATTGC | GCTGTGAT | HEG007 | AGCTAGTA | TGGAGCTT | SEG007 |
| GCTTCTAG | GATACTGC | AEG008 | GCTATTGC | GATACTGC | HEG008 | AGCTAGTA | GCTGTGAT | SEG008 |
| TAGCTCTG | TTACTGCA | AEG009 | AGTAACGG | TTACTGCA | HEG009 | TTACCTGA | ACTGTGTA | SEG009 |
| TAGCTCTG | ACTGTGTA | AEG010 | AGTAACGG | ACTGTGTA | HEG010 | TTACCTGA | GACTCTTA | SEG010 |
| TAGCTCTG | GACTCTTA | AEG011 | AGTAACGG | GACTCTTA | HEG011 | TTACCTGA | AACGTGGA | SEG011 |
| TAGCTCTG | TCGATCTC | AEG012 | AGTAACGG | AACGTGGA | HEG012 | TTACCTGA | TCGATCTC | SEG012 |
| TAGCTCTG | CTAGGCTT | AEG013 | AGTAACGG | TCGATCTC | HEG013 | TTACCTGA | CTAGGCTT | SEG013 |
| TAGCTCTG | TCGCTGAA | AEG014 | AGTAACGG | TCGCTGAA | HEG014 | TTACCTGA | TCGCTGAA | SEG014 |
| TAGCTCTG | AGGCTGTA | AEG015 | AGTAACGG | AGGCTGTA | HEG015 | TTACCTGA | AGGCTGTA | SEG015 |
| TAGCTCTG | TGACAGGT | AEG016 | AGTAACGG | TGACAGGT | HEG016 | TTACCTGA | TGACAGGT | SEG016 |
| GGTACTCA | CCTTAATG | AEG017 | ACTTCTGA | ATAGCTTG | HEG017 | TATACGAG | ATAGCTTG | SEG017 |
| GGTACTCA | CACATGCT | AEG018 | ACTTCTGA | CACATGCT | HEG018 | TATACGAG | CCTTAATG | SEG018 |
| GGTACTCA | TGTCATGC | AEG019 | ACTTCTGA | TGTCATGC | HEG019 | TATACGAG | CACATGCT | SEG019 |
| GGTACTCA | CGATAAGG | AEG020 | ACTTCTGA | CGATAAGG | HEG020 | TATACGAG | TGTCATGC | SEG020 |
| GGTACTCA | TTACGCGA | AEG021 | ACTTCTGA | TTACGCGA | HEG021 | TATACGAG | CGATAAGG | SEG021 |
| GGTACTCA | TGGAGCTT | AEG022 | ACTTCTGA | TGGAGCTT | HEG022 | TATACGAG | TTACGCGA | SEG022 |
| GGTACTCA | GCTGTGAT | AEG023 | ACTTCTGA | GCTGTGAT | HEG023 | TATACGAG | GCTGTGAT | SEG023 |
| GGTACTCA | GATACTGC | AEG024 | ACTTCTGA | GATACTGC | HEG024 | TATACGAG | GATACTGC | SEG024 |
| TGTAGGTC | TTACTGCA | AEG025 | ACTTAGCA | TTACTGCA | HEG025 | TATCAGTC | ACTGTGTA | SEG025 |
| TGTAGGTC | ACTGTGTA | AEG026 | ACTTAGCA | ACTGTGTA | HEG026 | TATCAGTC | GACTCTTA | SEG026 |
| TGTAGGTC | GACTCTTA | AEG027 | ACTTAGCA | AACGTGGA | HEG027 | TATCAGTC | AACGTGGA | SEG027 |
| TGTAGGTC | AACGTGGA | AEG028 | ACTTAGCA | TCGATCTC | HEG028 | TATCAGTC | TCGATCTC | SEG028 |
| TGTAGGTC | TCGATCTC | AEG029 | ACTTAGCA | CTAGGCTT | HEG029 | TATCAGTC | CTAGGCTT | SEG029 |
| TGTAGGTC | CTAGGCTT | AEG030 | ACTTAGCA | TCGCTGAA | HEG030 | TATCAGTC | TCGCTGAA | SEG030 |
| TGTAGGTC | AGGCTGTA | AEG031 | ACTTAGCA | AGGCTGTA | HEG031 | TATCAGTC | AGGCTGTA | SEG031 |
| TGTAGGTC | TGACAGGT | AEG032 | ACTTAGCA | TGACAGGT | HEG032 | TATCAGTC | TGACAGGT | SEG032 |
| TGCAACCA | CCTTAATG | AEG033 | AACTGTTC | CCTTAATG | HEG033 | TGACTTAG | CCTTAATG | SEG033 |
| TGCAACCA | CACATGCT | AEG034 | AACTGTTC | CACATGCT | HEG034 | TGACTTAG | CACATGCT | SEG034 |
| TGCAACCA | TGTCATGC | AEG035 | AACTGTTC | TGTCATGC | HEG035 | TGACTTAG | TGTCATGC | SEG035 |
| TGCAACCA | CGATAAGG | AEG036 | AACTGTTC | CGATAAGG | HEG036 | TGACTTAG | CGATAAGG | SEG036 |
| TGCAACCA | TTACGCGA | AEG037 | AACTGTTC | TTACGCGA | HEG037 | TGACTTAG | TTACGCGA | SEG037 |
| TGCAACCA | TGGAGCTT | AEG038 | AACTGTTC | TGGAGCTT | HEG038 | TGACTTAG | TGGAGCTT | SEG038 |
| TGCAACCA | GCTGTGAT | AEG039 | AACTGTTC | GCTGTGAT | HEG039 | TGACTTAG | GCTGTGAT | SEG039 |
| TGCAACCA | GATACTGC | AEG040 | AACTGTTC | GATACTGC | HEG040 | TGACTTAG | GATACTGC | SEG040 |
| CAATGCGA | ACTGTGTA | AEG041 | CATCTTGA | ACTGTGTA | HEG041 | TGTCAACA | TTACTGCA | SEG041 |
| CAATGCGA | GACTCTTA | AEG042 | CATCTTGA | GACTCTTA | HEG042 | TGTCAACA | ACTGTGTA | SEG042 |
| CAATGCGA | AACGTGGA | AEG043 | CATCTTGA | AACGTGGA | HEG043 | TGTCAACA | GACTCTTA | SEG043 |
| CAATGCGA | TCGATCTC | AEG044 | CATCTTGA | TCGATCTC | HEG044 | TGTCAACA | AACGTGGA | SEG044 |
| CAATGCGA | CTAGGCTT | AEG045 | CATCTTGA | CTAGGCTT | HEG045 | TGTCAACA | TCGATCTC | SEG045 |
| CAATGCGA | TCGCTGAA | AEG046 | CATCTTGA | TCGCTGAA | HEG046 | TGTCAACA | CTAGGCTT | SEG046 |
| CAATGCGA | AGGCTGTA | AEG047 | CATCTTGA | AGGCTGTA | HEG047 | TGTCAACA | TCGCTGAA | SEG047 |
| CAATGCGA | TGACAGGT | AEG048 | CATCTTGA | TGACAGGT | HEG048 | TGTCAACA | AGGCTGTA | SEG048 |
| CAATGCGA | TTACTGCA | AEG049 | AACTGTTC | ATAGCTTG | HEG049 | TGACTTAG | ATAGCTTG | SEG049 |
| GGTACTCA | ATAGCTTG | AEG050 | GCTATTGC | ATAGCTTG | HEG050 | TATCAGTC | TTACTGCA | SEG050 |
