## Supplementary material for "Distinct responses of oomycete plant parasites according to their lifestyle in a landscape-scale metabarcoding survey": Table S3

**Table S3**. References for the functional traits of the cercozoan genera identified in this study.

| Order | Genus | Lifestyle | Substrate | Detailed substrate | Above-, below-ground | Usual name | References | Remarks |
|---|---|---|---|---|---|---|---|---|
| Albuginales | Pustula | obligate biotroph | Plantae | NA | | white blister rust | 1 | |
| Haptoglossales | Haptoglossa | obligate biotroph | Metazoa | Nematoda, Rotifera | belowground | | 1 | |
| Leptomitales | Apodachlya | saprotroph | NA | NA | | water moulds | 1 | |
| Leptomitales | Blastulidium | hemibiotroph | Metazoa | Cladocera | aquatic | | 1 | |
| Peronosporales | Globisporangium | obligate biotroph | Plantae | generalist | infection below ground | seed rots and dampingoff | 1 | 1 |
| Peronosporales | Hyaloperonospora | obligate biotroph | Plantae | Brassicaceae | | Brassicolous downy mildews | 2 | |
| Peronosporales | Lagena | obligate biotroph | Plantae | Poaceae | | | 3 | |
| Peronosporales | Lagenidium | hemibiotroph | Metazoa | NA | | | 4 | |
| Peronosporales | Myzocytiopsis | hemibiotroph | Metazoa | Nematoda | belowground | | 1 | |
| Peronosporales | Paralagenidium | hemibiotroph | Metazoa | NA | | | 4 | |
| Peronosporales | Peronospora | obligate biotroph | Plantae | specilialist | | downy mildews | 2 | 2 |
| Peronosporales | Peronosporaceae | NA | NA | NA | | | | |
| Peronosporales | Phytophthora | hemibiotroph | Plantae | NA | | | 5 | 3 |
| Peronosporales | Phytopythium | saprotroph | Plantae | generalist | belowground | saprotrophic Peronosporales | 6 | |
| Peronosporales | Plasmopara | obligate biotroph | Plantae | specilialist | | downy mildew | 7 | 2 |
| Peronosporales | Plasmoverna | obligate biotroph | Plantae | Ranunculaceae | | | 7 | |
| Peronosporales | Protobremia | obligate biotroph | Plantae | NA | | | 7 | |
| Peronosporales | Pythiaceae | NA | NA | NA | | | | |
| Peronosporales | Pythiogeton | NA | NA | NA | | | 8 | |
| Peronosporales | Pythium | hemibiotroph | Plantae | NA | belowground | seed rots and dampingoff | 5 | 4 |
| Saprolegniales | Achlya | hemibiotroph | NA | NA | aquatic | water moulds | 9 | |
| Saprolegniales | Aphanomyces | NA | NA | NA | | | 1 | |
| Saprolegniales | Aplanopsis | saprotroph | NA | NA | belowground, aquatic | | 10 | |
| Saprolegniales | Atkinsiella | hemibiotroph | Metazoa | Crustacea | aquatic | | 1 | |
| Saprolegniales | Brevilegnia | saprotroph | NA | NA | belowground, aquatic | water moulds | 1 | |
| Saprolegniales | Dictyuchus | saprotroph | NA | NA | belowground, aquatic | water moulds | 1 | |
| Saprolegniales | Leptolegnia | saprotroph | Metazoa | exuviae of insects | belowground, aquatic | water moulds | 11 | |
| Saprolegniales | Plectospira | saprotroph | Plantae | NA | belowground, aquatic | water moulds | 11 | |
| Saprolegniales | Pythiopsis | NA | NA | NA | | | | |
| Saprolegniales | Saprolegnia | hemibiotroph | Metazoa | fish and their eggs, moll | belowground, aquatic | water moulds | 11 | |
| Saprolegniales | Saprolegniaceae | NA | NA | NA | | water moulds | 11 | |
| Saprolegniales | Thraustotheca | saprotroph | NA | NA | belowground, aquatic | water moulds | 11 | |

### *References:*

7. Plant parasites of Europe, https://bladmineerders.nl/

8. Huang J-H, Chen C-Y, Lin Y-S, Ann P-J, Huang H-C & Chung W-H (2013) Six new species of Pythiogeton in Taiwan, with an account of the molecular phylogeny of this genus. Mycoscience 54: 130-147.

9. Singhal R, Jeet S & Davies R (1987) Experimental transmission of Saprolegnia and Achlya to fish. Aquaculture 64: 1-7.

10. Spencer MA, Vick MC & Dick MW (2002) Revision of Aplanopsis, Pythiopsis, and ‘subcentric’ Achlya species (Saprolegniaceae) using 18S rDNA and morphological data. Mycological Research 106: 549560.

11. Johnson, T.W., Seymour, R.L., & Padgett, D.E. (2002). Biology and systematics of the Saprolegniaceae. http://dl.uncw.edu/digilib/Biology/Fungi/Taxonomy%20and%20Systematics/Padgett%20Book/


### *Remarks:*

1. Genus recently separated from *Pythium* . *Phythium sylvaticum* is now *Globisporangium*
2. Ususally not generalist: different species infect specific plant families
3. Only few species of *Phyophthora* are saprotrophs
4. Only *P. insidiosum* infects mammals (quine phycomycosis)
