## Supplementary material for "Distinct responses of oomycete plant parasites according to their lifestyle in a landscape-scale metabarcoding survey": Table S5

### Table S5. Environmental parameters from the 150 forest study sites and two years of collection.

| Reference dataset at https://www.bexis.uni-jena.de/ | 14447 22246 | 14446 23846 | 14446 23846 | 14446 23846 | 14446 23846 | 14446 23846 | | 14686 | | 16466 | 10580 | 17706 | 17706 | 17706 |
|---|---|---|---|---|---|---|---|---|---|---|---|---|---|---|
| Sites (AE=Alb, HE=Hainich, SE=Schorfheide, 11/17:year of collection | pH | Total_C g/kg soil | Inorganic_C g/kg | Organic_C g/kg | Total_N g/kg | CN_ratio | Clay | Silt | Sand | Intensity_management[1] | soil_type | main_tree_species | Pure_or_mixed[2] | Developmental_stage[3] |
| AEF008_11 | 6.08 | 62.18 | 1.77 | 60.41 | 4.28 | 14.11 | 492 | 466 | 42 | 0.0000 | Cambisol | Fs | 3_mixed | 50_100 |
| AEF049_11 | 5.86 | 75.33 | 0.66 | 74.67 | 5.90 | 12.65 | 645 | 234 | 121 | 0.7165 | Leptosol | Fs | 3_mixed | 50_100 |
| AEF049_17 | 6.31 | 77.28 | 0.68 | 76.60 | 5.79 | 13.23 | 645 | 234 | 121 | 0.7165 | Leptosol | Fs | 3_mixed | 50_100 |
| AEF048_11 | 5.40 | 59.08 | 0.41 | 58.67 | 4.82 | 12.18 | 464 | 508 | 28 | 1.5792 | Leptosol | Fs | 3_mixed | 30_50 |
| AEF048_17 | 5.75 | 69.75 | 0.48 | 69.28 | 5.25 | 13.20 | 464 | 508 | 28 | 1.5792 | Leptosol | Fs | 3_mixed | 30_50 |
| HEF039_11 | 4.14 | 31.88 | 0.25 | 31.63 | 2.50 | 12.64 | 330 | 615 | 58 | 0.0928 | Luvisol | Fs | 3_mixed | 50_100 |
| HEF039_17 | 4.52 | 38.87 | 0.00 | 38.87 | 3.14 | 12.37 | 330 | 615 | 58 | 0.0928 | Luvisol | Fs | 3_mixed | 50_100 |
| AEF008_17 | 6.43 | 60.06 | 1.17 | 58.89 | 4.17 | 14.11 | 492 | 466 | 42 | 0.0000 | Cambisol | Fs | 3_mixed | 50_100 |
| SEF032_11 | 3.21 | 28.55 | 0.00 | 28.55 | 1.44 | 19.78 | 51 | 141 | 808 | 1.7734 | Cambisol | Ps | 3_mixed | 50_100 |
| SEF032_17 | 3.53 | 22.63 | 0.00 | 22.63 | 1.14 | 19.92 | 51 | 141 | 808 | 1.7734 | Cambisol | Ps | 3_mixed | 50_100 |
| SEF048_11 | 3.50 | 19.33 | 0.00 | 19.33 | 1.17 | 16.46 | 85 | 190 | 725 | 0.4675 | Cambisol | Fs | 3_mixed | 50_100 |
| SEF048_17 | 3.67 | 23.36 | 0.00 | 23.36 | 1.26 | 18.59 | 85 | 190 | 725 | 0.4675 | Cambisol | Fs | 3_mixed | 50_100 |
| SEF024_11 | 3.44 | 18.17 | 0.00 | 18.17 | 1.15 | 15.80 | 46 | 142 | 812 | 0.6409 | Cambisol | Qs | 3_mixed | 50_100 |
| SEF024_17 | 3.66 | 10.48 | 0.00 | 10.48 | 0.68 | 15.31 | 46 | 142 | 812 | 0.6409 | Cambisol | Qs | 3_mixed | 50_100 |
| AEF015_11 | 6.23 | 94.53 | 0.98 | 93.55 | 6.80 | 13.76 | 653 | 277 | 70 | 2.8220 | Leptosol | Fs | Fs_oHS_mixed | 0-15 |
| AEF019_11 | 4.80 | 60.17 | 0.28 | 59.89 | 4.96 | 12.08 | 471 | 503 | 28 | 0.6589 | Cambisol | Fs | Fs_oHS_mixed | 50_100 |
| AEF025_11 | 4.70 | 39.19 | 0.24 | 38.95 | 3.02 | 12.90 | 383 | 506 | 111 | 1.7803 | Cambisol | Fs | Fs_oHS_mixed | 0-15 |
| AEF027_11 | 4.49 | 58.50 | 0.41 | 58.09 | 4.21 | 13.80 | 518 | 352 | 130 | 2.6402 | Leptosol | Fs | Fs_oHS_mixed | 15_30 |
| AEF028_11 | 4.90 | 61.57 | 0.29 | 61.28 | 4.80 | 12.75 | 474 | 474 | 52 | 1.2266 | Cambisol | Fs | Fs_oHS_mixed | 30_50 |
| AEF030_11 | 5.94 | 86.82 | 0.89 | 85.92 | 6.84 | 12.55 | 700 | 263 | 37 | 1.1701 | Cambisol | Fs | Fs_oHS_mixed | 30_50 |
| AEF035_11 | 5.11 | 83.17 | 0.42 | 82.75 | 5.77 | 14.34 | 587 | 362 | 51 | 2.6049 | Leptosol | Fs | Fs_oHS_mixed | 0-15 |
| AEF036_11 | 5.85 | 65.20 | 0.56 | 64.63 | 5.36 | 12.06 | 407 | 541 | 52 | 1.9237 | Leptosol | Fs | Fs_oHS_mixed | 0-15 |
| AEF037_11 | 5.40 | 59.22 | 0.38 | 58.85 | 4.73 | 12.44 | 553 | 374 | 73 | 0.9939 | Leptosol | Fs | Fs_oHS_mixed | 0-15 |
| AEF044_11 | 5.89 | 61.59 | 0.40 | 61.19 | 4.97 | 12.31 | 429 | 528 | 43 | 0.7599 | Leptosol | Fs | Fs_oHS_mixed | 0-15 |
| AEF046_11 | 5.07 | 53.51 | 0.28 | 53.22 | 4.14 | 12.85 | 546 | 425 | 27 | 1.1088 | Leptosol | Fs | Fs_oHS_mixed | 30_50 |
| HEF004_11 | 6.00 | 47.07 | 0.46 | 46.61 | 3.68 | 12.66 | 387 | 562 | 55 | 1.8905 | Luvisol | Fs | Fs_oHS_mixed | 0-15 |
| HEF010_11 | 4.86 | 50.88 | 0.33 | 50.55 | 4.04 | 12.50 | 447 | 503 | 53 | 0.0641 | Stagnosol | Fs | Fs_oHS_mixed | 50_100 |
| HEF011_11 | 4.53 | 37.62 | 0.30 | 37.32 | 3.02 | 12.35 | 414 | 551 | 36 | 0.5177 | Luvisol | Fs | Fs_oHS_mixed | 50_100 |
| HEF014_11 | 4.54 | 44.47 | 0.39 | 44.08 | 3.34 | 13.21 | 448 | 504 | 47 | 1.6478 | Luvisol | Fs | Fs_oHS_mixed | 0-15 |
| HEF015_11 | 3.91 | 20.86 | 0.00 | 20.86 | 1.29 | 16.20 | 150 | 785 | 66 | 1.8782 | Luvisol | Fs | Fs_oHS_mixed | 0-15 |
| HEF017_11 | 3.86 | 27.28 | 0.00 | 27.28 | 1.79 | 15.24 | 183 | 725 | 96 | 1.1114 | Luvisol | Fs | Fs_oHS_mixed | 15_30 |
| HEF018_11 | 4.79 | 35.14 | 0.23 | 34.91 | 2.69 | 12.99 | 272 | 684 | 41 | 1.3244 | Stagnosol | Fs | Fs_oHS_mixed | 15_30 |
| HEF019_11 | 4.57 | 31.94 | 0.28 | 31.66 | 2.78 | 11.40 | 310 | 636 | 52 | 1.0176 | Luvisol | Fs | Fs_oHS_mixed | 30_50 |
| HEF034_11 | 4.62 | 38.01 | 0.25 | 37.75 | 3.19 | 11.84 | 380 | 571 | 52 | 0.0000 | Luvisol | Fs | Fs_oHS_mixed | 50_100 |
| HEF035_11 | 4.50 | 36.45 | 0.27 | 36.18 | 2.91 | 12.44 | 396 | 551 | 55 | 0.8610 | Luvisol | Fs | Fs_oHS_mixed | 50_100 |
| HEF036_11 | 4.56 | 37.76 | 0.24 | 37.52 | 2.92 | 12.84 | 356 | 558 | 84 | 0.3653 | Luvisol | Fs | Fs_oHS_mixed | 50_100 |
| HEF038_11 | 5.15 | 53.04 | 0.37 | 52.67 | 4.04 | 13.04 | 511 | 440 | 54 | 0.0480 | Luvisol | Fs | Fs_oHS_mixed | 50_100 |
| HEF040_11 | 5.47 | 48.95 | 0.44 | 48.51 | 4.02 | 12.05 | 406 | 563 | 33 | 0.0230 | Luvisol | Fs | Fs_oHS_mixed | 50_100 |
| HEF041_11 | 4.26 | 24.25 | 0.00 | 24.25 | 1.94 | 12.52 | 210 | 754 | 34 | 0.3200 | Luvisol | Fs | Fs_oHS_mixed | 50_100 |
| HEF043_11 | 6.35 | 50.98 | 1.04 | 49.94 | 3.89 | 12.83 | 337 | 635 | 30 | 1.8559 | Stagnosol | Fs | Fs_oHS_mixed | 0-15 |
| HEF044_11 | 6.27 | 46.69 | 0.80 | 45.89 | 3.42 | 13.42 | 373 | 586 | 43 | 1.7997 | Stagnosol | Fs | Fs_oHS_mixed | 0-15 |
| AEF015_17 | 6.37 | 65.02 | 1.01 | 64.01 | 4.80 | 13.34 | 653 | 277 | 70 | 2.8220 | Leptosol | Fs | Fs_oHS_mixed | 0-15 |
| AEF019_17 | 5.07 | 58.13 | 0.47 | 57.66 | 4.74 | 12.16 | 471 | 503 | 28 | 0.6589 | Cambisol | Fs | Fs_oHS_mixed | 50_100 |

| | | | | | | | | | | | | | | |
|---|---|---|---|---|---|---|---|---|---|---|---|---|---|---|
| AEF025_17 | 5.13 | 41.73 | 0.00 | 41.73 | 3.26 | 12.79 | 383 | 506 | 111 | 1.7803 | Cambisol | Fs | Fs_oHS_mixed | 0-15 |
| AEF027_17 | 4.58 | 50.21 | 0.00 | 50.21 | 3.72 | 13.50 | 518 | 352 | 130 | 2.6402 | Leptosol | Fs | Fs_oHS_mixed | 15_30 |
| AEF028_17 | 4.73 | 54.18 | 0.31 | 53.87 | 4.12 | 13.06 | 474 | 474 | 52 | 1.2266 | Cambisol | Fs | Fs_oHS_mixed | 30_50 |
| AEF030_17 | 5.82 | 84.03 | 0.91 | 83.12 | 6.60 | 12.59 | 700 | 263 | 37 | 1.1701 | Cambisol | Fs | Fs_oHS_mixed | 30_50 |
| AEF035_17 | 5.48 | 79.91 | 0.48 | 79.43 | 5.61 | 14.15 | 587 | 362 | 51 | 2.6049 | Leptosol | Fs | Fs_oHS_mixed | 0-15 |
| AEF036_17 | 5.98 | 62.41 | 0.49 | 61.92 | 4.82 | 12.86 | 407 | 541 | 52 | 1.9237 | Leptosol | Fs | Fs_oHS_mixed | 0-15 |
| AEF037_17 | 5.22 | 58.76 | 0.35 | 58.41 | 4.58 | 12.75 | 553 | 374 | 73 | 0.9939 | Leptosol | Fs | Fs_oHS_mixed | 0-15 |
| AEF044_17 | 6.04 | 63.60 | 0.45 | 63.16 | 4.76 | 13.27 | 429 | 528 | 43 | 0.7599 | Leptosol | Fs | Fs_oHS_mixed | 0-15 |
| AEF046_17 | 5.45 | 55.54 | 0.31 | 55.22 | 4.10 | 13.46 | 546 | 425 | 27 | 1.1088 | Leptosol | Fs | Fs_oHS_mixed | 30_50 |
| HEF004_17 | 6.14 | 63.48 | 0.53 | 62.95 | 4.77 | 13.19 | 387 | 562 | 55 | 1.8905 | Luvisol | Fs | Fs_oHS_mixed | 0-15 |
| HEF010_17 | 4.93 | 52.41 | 0.00 | 52.41 | 4.03 | 13.00 | 447 | 503 | 53 | 0.0641 | Stagnosol | Fs | Fs_oHS_mixed | 50_100 |
| HEF011_17 | 4.87 | 50.69 | 0.00 | 50.69 | 3.83 | 13.22 | 414 | 551 | 36 | 0.5177 | Luvisol | Fs | Fs_oHS_mixed | 50_100 |
| HEF014_17 | 5.05 | 60.73 | 0.49 | 60.23 | 4.35 | 13.86 | 448 | 504 | 47 | 1.6478 | Luvisol | Fs | Fs_oHS_mixed | 0-15 |
| HEF015_17 | 3.99 | 28.14 | 0.00 | 28.14 | 1.89 | 14.89 | 150 | 785 | 66 | 1.8782 | Luvisol | Fs | Fs_oHS_mixed | 0-15 |
| HEF017_17 | 3.87 | 38.17 | 0.00 | 38.17 | 2.55 | 14.96 | 183 | 725 | 96 | 1.1114 | Luvisol | Fs | Fs_oHS_mixed | 15_30 |
| HEF018_17 | 5.55 | 42.56 | 0.55 | 42.01 | 3.05 | 13.78 | 272 | 684 | 41 | 1.3244 | Stagnosol | Fs | Fs_oHS_mixed | 15_30 |
| HEF019_17 | 4.61 | 33.75 | 0.00 | 33.75 | 2.98 | 11.33 | 310 | 636 | 52 | 1.0176 | Luvisol | Fs | Fs_oHS_mixed | 30_50 |
| HEF034_17 | 4.66 | 40.10 | 0.00 | 40.10 | 3.25 | 12.35 | 380 | 571 | 52 | 0.0000 | Luvisol | Fs | Fs_oHS_mixed | 50_100 |
| HEF035_17 | 4.44 | 40.64 | 0.00 | 40.64 | 3.33 | 12.19 | 396 | 551 | 55 | 0.8610 | Luvisol | Fs | Fs_oHS_mixed | 50_100 |
| HEF036_17 | 4.71 | 44.54 | 0.00 | 44.54 | 3.25 | 13.71 | 356 | 558 | 84 | 0.3653 | Luvisol | Fs | Fs_oHS_mixed | 50_100 |
| HEF038_17 | 5.42 | 70.91 | 0.49 | 70.42 | 5.24 | 13.45 | 511 | 440 | 54 | 0.0480 | Luvisol | Fs | Fs_oHS_mixed | 50_100 |
| HEF040_17 | 5.43 | 53.10 | 0.00 | 53.10 | 3.94 | 13.47 | 406 | 563 | 33 | 0.0230 | Luvisol | Fs | Fs_oHS_mixed | 50_100 |
| HEF041_17 | 4.55 | 31.89 | 0.00 | 31.89 | 2.51 | 12.70 | 210 | 754 | 34 | 0.3200 | Luvisol | Fs | Fs_oHS_mixed | 50_100 |
| HEF043_17 | 6.72 | 69.24 | 2.60 | 66.64 | 4.86 | 13.71 | 337 | 635 | 30 | 1.8559 | Stagnosol | Fs | Fs_oHS_mixed | 0-15 |
| HEF044_17 | 5.36 | 56.58 | 0.48 | 56.10 | 3.93 | 14.27 | 373 | 586 | 43 | 1.7997 | Stagnosol | Fs | Fs_oHS_mixed | 0-15 |
| AEF007_11 | 4.60 | 62.64 | 0.22 | 62.42 | 4.78 | 13.05 | 548 | 362 | 90 | 1.0285 | Leptosol | Fs | Fs_Pa_mixed | 50_100 |
| AEF047_11 | 5.07 | 67.28 | 0.39 | 66.88 | 5.21 | 12.84 | 531 | 404 | 65 | 1.1534 | Cambisol | Fs | Fs_Pa_mixed | 30_50 |
| AEF007_17 | 5.01 | 62.89 | 0.39 | 62.50 | 4.85 | 12.89 | 548 | 362 | 90 | 1.0285 | Leptosol | Fs | Fs_Pa_mixed | 50_100 |
| AEF047_17 | 5.15 | 60.63 | 0.29 | 60.33 | 4.63 | 13.03 | 531 | 404 | 65 | 1.1534 | Cambisol | Fs | Fs_Pa_mixed | 30_50 |
| SEF030_11 | 3.12 | 26.81 | 0.00 | 26.81 | 1.16 | 23.06 | 50 | 172 | 778 | 1.3926 | Cambisol | Ps | Fs_Ps_mixed | 50_100 |
| SEF034_11 | 3.22 | 16.44 | 0.00 | 16.44 | 0.74 | 22.22 | 42 | 69 | 889 | 1.7943 | Albeluvisol | Ps | Fs_Ps_mixed | 50_100 |
| SEF030_17 | 3.44 | 19.29 | 0.00 | 19.29 | 0.94 | 20.58 | 50 | 172 | 778 | 1.3926 | Cambisol | Ps | Fs_Ps_mixed | 50_100 |
| SEF034_17 | 3.46 | 16.91 | 0.00 | 16.91 | 0.75 | 22.57 | 42 | 69 | 889 | 1.7943 | Albeluvisol | Ps | Fs_Ps_mixed | 50_100 |
| AEF004_11 | 6.20 | 71.22 | 0.90 | 70.32 | 5.51 | 12.75 | 487 | 482 | 32 | 1.5633 | Cambisol | Fs | Fs_pure | 15_30 |
| AEF005_11 | 4.51 | 48.61 | 0.21 | 48.40 | 3.73 | 12.98 | 404 | 579 | 18 | 0.9392 | Cambisol | Fs | Fs_pure | 50_100 |
| AEF006_11 | 4.90 | 39.78 | 0.19 | 39.58 | 3.21 | 12.32 | 415 | 494 | 91 | 1.1882 | Cambisol | Fs | Fs_pure | 30_50 |
| AEF009_11 | 5.90 | 58.64 | 0.83 | 57.81 | 4.05 | 14.29 | 693 | 289 | 18 | 0.4853 | Leptosol | Fs | Fs_pure | 50_100 |
| AEF016_11 | 6.20 | 60.63 | 0.67 | 59.96 | 4.34 | 13.81 | 521 | 335 | 144 | 1.0864 | Cambisol | Fs | Fs_pure | 0-15 |
| AEF017_11 | 6.44 | 53.26 | 1.55 | 51.72 | 4.13 | 12.50 | 428 | 540 | 32 | 1.2564 | Cambisol | Fs | Fs_pure | 15_30 |
| AEF018_11 | 4.62 | 35.96 | 0.22 | 35.73 | 2.81 | 12.71 | 313 | 655 | 34 | 0.3935 | Cambisol | Fs | Fs_pure | 50_100 |
| AEF020_11 | 6.64 | 59.60 | 1.28 | 58.32 | 4.80 | 12.14 | 437 | 533 | 30 | 1.0784 | Cambisol | Fs | Fs_pure | 50_100 |
| AEF022_11 | 6.25 | 59.29 | 0.56 | 58.73 | 4.41 | 13.31 | 542 | 409 | 49 | 0.7178 | Cambisol | Fs | Fs_pure | 30_50 |
| AEF023_11 | 5.48 | 58.19 | 0.38 | 57.81 | 4.65 | 12.42 | 500 | 486 | 15 | 1.2499 | Cambisol | Fs | Fs_pure | 50_100 |
| AEF038_11 | 6.17 | 56.64 | 0.63 | 56.01 | 4.17 | 13.42 | 502 | 387 | 111 | 1.5666 | Leptosol | Fs | Fs_pure | 15_30 |
| AEF039_11 | 5.75 | 59.61 | 0.53 | 59.07 | 4.51 | 13.10 | 432 | 530 | 38 | 1.2416 | Leptosol | Fs | Fs_pure | 15_30 |
| AEF040_11 | 5.01 | 75.70 | 0.33 | 75.36 | 6.58 | 11.46 | 610 | 347 | 44 | 0.9907 | Leptosol | Fs | Fs_pure | 50_100 |
| AEF041_11 | 5.12 | 60.31 | 0.41 | 59.91 | 4.87 | 12.30 | 475 | 449 | 76 | 1.0057 | Leptosol | Fs | Fs_pure | 30_50 |
| AEF042_11 | 5.88 | 65.27 | 0.55 | 64.72 | 5.15 | 12.57 | 504 | 386 | 110 | 1.2601 | Leptosol | Fs | Fs_pure | 30_50 |
| AEF043_11 | 4.80 | 37.01 | 0.22 | 36.79 | 3.32 | 11.06 | 386 | 577 | 39 | 1.0001 | Leptosol | Fs | Fs_pure | 50_100 |
| AEF050_11 | 5.87 | 100.40 | 0.90 | 99.55 | 7.55 | 13.19 | 543 | 429 | 28 | 0.9005 | Leptosol | Fs | Fs_pure | 50_100 |
| HEF005_11 | 4.99 | 43.84 | 0.61 | 43.22 | 3.42 | 12.64 | 460 | 485 | 59 | 0.9556 | Luvisol | Fs | Fs_pure | 30_50 |
| HEF006_11 | 4.15 | 22.88 | 0.22 | 22.66 | 1.82 | 12.46 | 218 | 709 | 74 | 0.7467 | Luvisol | Fs | Fs_pure | 50_100 |
| HEF007_11 | 4.10 | 28.98 | 0.22 | 28.76 | 2.06 | 13.95 | 197 | 714 | 88 | 0.6948 | Luvisol | Fs | Fs_pure | 50_100 |
| HEF008_11 | 5.56 | 27.87 | 0.28 | 27.59 | 1.98 | 13.93 | 227 | 716 | 58 | 1.1562 | Luvisol | Fs | Fs_pure | 50_100 |

| | | | | | | | | | | | | | | |
|---|---|---|---|---|---|---|---|---|---|---|---|---|---|---|
| HEF009_11 | 4.12 | 30.24 | 0.28 | 29.97 | 2.34 | 12.78 | 287 | 622 | 92 | 1.2574 | Luvisol | Fs | Fs_pure | 50_100 |
| HEF012_11 | 4.05 | 20.93 | 0.00 | 20.93 | 1.56 | 13.41 | 168 | 713 | 117 | 0.0000 | Luvisol | Fs | Fs_pure | 50_100 |
| HEF020_11 | 5.97 | 32.59 | 0.65 | 31.94 | 2.46 | 12.97 | 253 | 708 | 40 | 0.9926 | Luvisol | Fs | Fs_pure | 30_50 |
| HEF021_11 | 4.71 | 29.44 | 0.31 | 29.13 | 2.34 | 12.46 | 268 | 672 | 61 | 0.8144 | Luvisol | Fs | Fs_pure | 50_100 |
| HEF022_11 | 4.63 | 23.33 | 0.00 | 23.33 | 1.67 | 13.94 | 184 | 747 | 68 | 0.6386 | Luvisol | Fs | Fs_pure | 50_100 |
| HEF023_11 | 4.53 | 34.49 | 0.22 | 34.27 | 2.73 | 12.56 | 279 | 662 | 56 | 1.3847 | Luvisol | Fs | Fs_pure | 50_100 |
| HEF024_11 | 4.05 | 22.31 | 0.44 | 21.87 | 1.59 | 13.73 | 162 | 793 | 44 | 0.8871 | Luvisol | Fs | Fs_pure | 50_100 |
| HEF025_11 | 4.58 | 39.61 | 0.25 | 39.35 | 3.09 | 12.73 | 325 | 624 | 53 | 1.2635 | Luvisol | Fs | Fs_pure | 50_100 |
| HEF026_11 | 4.46 | 24.42 | 0.00 | 24.42 | 1.63 | 14.95 | 150 | 796 | 54 | 1.1968 | Luvisol | Fs | Fs_pure | 50_100 |
| HEF027_11 | 6.37 | 50.10 | 0.68 | 49.42 | 3.82 | 12.93 | 291 | 643 | 67 | 0.7979 | Luvisol | Fs | Fs_pure | 50_100 |
| HEF028_11 | 6.29 | 44.68 | 0.53 | 44.15 | 3.20 | 13.80 | 306 | 634 | 60 | 1.4438 | Stagnosol | Fs | Fs_pure | 50_100 |
| HEF029_11 | 3.86 | 25.90 | 0.00 | 25.90 | 1.86 | 13.94 | 227 | 719 | 55 | 0.5728 | Luvisol | Fs | Fs_pure | 50_100 |
| HEF030_11 | 3.86 | 33.78 | 0.29 | 33.48 | 2.66 | 12.57 | 358 | 595 | 48 | 0.7626 | Luvisol | Fs | Fs_pure | 50_100 |
| HEF031_11 | 3.88 | 33.36 | 0.32 | 33.04 | 2.62 | 12.62 | 377 | 573 | 52 | 1.0788 | Luvisol | Fs | Fs_pure | 50_100 |
| HEF032_11 | 3.92 | 36.27 | 0.25 | 36.02 | 2.81 | 12.84 | 345 | 593 | 64 | 1.1500 | Luvisol | Fs | Fs_pure | 50_100 |
| HEF033_11 | 4.45 | 27.93 | 0.23 | 27.70 | 1.95 | 14.17 | 227 | 706 | 68 | 1.1054 | Luvisol | Fs | Fs_pure | 50_100 |
| HEF037_11 | 4.50 | 31.80 | 0.23 | 31.57 | 2.50 | 12.61 | 265 | 661 | 78 | 0.0000 | Stagnosol | Fs | Fs_pure | 50_100 |
| HEF042_11 | 3.97 | 23.41 | 0.00 | 23.41 | 1.63 | 14.34 | 184 | 760 | 60 | 0.5400 | Stagnosol | Fs | Fs_pure | 50_100 |
| HEF045_11 | 7.10 | 67.36 | 9.94 | 57.42 | 4.60 | 12.50 | 88 | 854 | 60 | 1.0203 | Luvisol | Fs | Fs_pure | 15_30 |
| HEF046_11 | 3.92 | 37.14 | 0.22 | 36.92 | 2.91 | 12.70 | 387 | 570 | 45 | 1.0506 | Luvisol | Fs | Fs_pure | 30_50 |
| HEF047_11 | 4.79 | 32.51 | 0.00 | 32.51 | 2.43 | 13.35 | 323 | 632 | 46 | 0.8971 | Stagnosol | Fs | Fs_pure | 50_100 |
| HEF048_11 | 4.13 | 26.65 | 0.00 | 26.65 | 2.02 | 13.21 | 273 | 687 | 44 | 0.8371 | Stagnosol | Fs | Fs_pure | 50_100 |
| HEF049_11 | 3.89 | 23.82 | 0.00 | 23.82 | 1.61 | 14.83 | 223 | 732 | 48 | 1.2523 | Stagnosol | Fs | Fs_pure | 50_100 |
| HEF050_11 | 4.75 | 33.56 | 0.23 | 33.32 | 2.80 | 11.91 | 349 | 606 | 46 | 0.4231 | Stagnosol | Fs | Fs_pure | 50_100 |
| SEF005_11 | 3.18 | 25.04 | 0.00 | 25.04 | 1.29 | 19.37 | 19 | 17 | 964 | 0.6890 | Cambisol | Fs | Fs_pure | 50_100 |
| SEF006_11 | 3.45 | 25.87 | 0.00 | 25.87 | 1.49 | 17.39 | 49 | 77 | 874 | 1.4018 | Cambisol | Fs | Fs_pure | 50_100 |
| SEF007_11 | 3.49 | 18.65 | 0.00 | 18.65 | 1.15 | 16.29 | 16 | 124 | 860 | 0.0855 | Cambisol | Fs | Fs_pure | 50_100 |
| SEF008_11 | 3.28 | 27.09 | 0.00 | 27.09 | 1.64 | 16.54 | 47 | 198 | 755 | 0.1873 | Albeluvisol | Fs | Fs_pure | 50_100 |
| SEF009_11 | 3.27 | 20.01 | 0.00 | 20.01 | 1.04 | 19.18 | 0 | 58 | 942 | 0.6459 | Cambisol | Fs | Fs_pure | 50_100 |
| SEF035_11 | 3.43 | 20.64 | 0.00 | 20.64 | 1.10 | 18.82 | 65 | 74 | 861 | 1.0252 | Cambisol | Fs | Fs_pure | 50_100 |
| SEF036_11 | 3.21 | 30.36 | 0.00 | 30.36 | 1.76 | 17.26 | 64 | 88 | 848 | 1.2034 | Cambisol | Fs | Fs_pure | 30_50 |
| SEF039_11 | 3.46 | 18.94 | 0.00 | 18.94 | 1.07 | 17.74 | 52 | 98 | 850 | 1.0994 | Cambisol | Fs | Fs_pure | 50_100 |
| SEF040_11 | 3.56 | 18.97 | 0.00 | 18.97 | 1.07 | 17.73 | 63 | 71 | 866 | 1.3649 | Cambisol | Fs | Fs_pure | 50_100 |
| SEF041_11 | 3.57 | 17.59 | 0.00 | 17.59 | 0.93 | 18.96 | 50 | 49 | 905 | 1.1019 | Cambisol | Fs | Fs_pure | 50_100 |
| SEF042_11 | 3.55 | 23.20 | 0.00 | 23.20 | 1.38 | 16.85 | 74 | 247 | 679 | 0.7863 | Regosol | Fs | Fs_pure | 50_100 |
| SEF043_11 | 3.48 | 18.41 | 0.00 | 18.41 | 1.12 | 16.49 | 69 | 121 | 810 | 1.0317 | Cambisol | Fs | Fs_pure | 50_100 |
| SEF045_11 | 3.50 | 16.51 | 0.00 | 16.51 | 0.92 | 17.90 | 48 | 75 | 881 | 0.3413 | Albeluvisol | Fs | Fs_pure | 50_100 |
| SEF046_11 | 3.30 | 21.20 | 0.00 | 21.20 | 1.34 | 15.87 | 63 | 132 | 805 | 0.2069 | Albeluvisol | Fs | Fs_pure | 50_100 |
| SEF047_11 | 3.21 | 23.50 | 0.00 | 23.50 | 1.33 | 17.62 | 49 | 18 | 933 | 0.0156 | Cambisol | Fs | Fs_pure | 50_100 |
| SEF049_11 | 3.24 | 21.39 | 0.00 | 21.39 | 1.15 | 18.52 | 68 | 74 | 858 | 1.0371 | Cambisol | Fs | Fs_pure | 50_100 |
| SEF050_11 | 3.41 | 16.73 | 0.00 | 16.73 | 1.04 | 16.08 | 79 | 99 | 822 | 1.1073 | Cambisol | Fs | Fs_pure | 50_100 |
| AEF004_17 | 6.76 | 77.36 | 3.85 | 73.51 | 6.00 | 12.25 | 487 | 482 | 32 | 1.5633 | Cambisol | Fs | Fs_pure | 15_30 |
| AEF005_17 | 4.45 | 46.44 | 0.33 | 46.11 | 3.58 | 12.86 | 404 | 579 | 18 | 0.9392 | Cambisol | Fs | Fs_pure | 50_100 |
| AEF006_17 | 5.59 | 47.95 | 0.44 | 47.51 | 3.76 | 12.63 | 415 | 494 | 91 | 1.1882 | Cambisol | Fs | Fs_pure | 30_50 |
| AEF009_17 | 6.12 | 61.25 | 0.66 | 60.59 | 4.27 | 14.20 | 693 | 289 | 18 | 0.4853 | Leptosol | Fs | Fs_pure | 50_100 |
| AEF016_17 | 6.37 | 67.50 | 0.90 | 66.61 | 4.90 | 13.60 | 521 | 335 | 144 | 1.0864 | Cambisol | Fs | Fs_pure | 0-15 |
| AEF017_17 | 6.53 | 52.02 | 1.10 | 50.92 | 4.14 | 12.31 | 428 | 540 | 32 | 1.2564 | Cambisol | Fs | Fs_pure | 15_30 |
| AEF018_17 | 4.69 | 43.55 | 0.29 | 43.25 | 3.36 | 12.88 | 313 | 655 | 34 | 0.3935 | Cambisol | Fs | Fs_pure | 50_100 |
| AEF020_17 | 6.60 | 72.16 | 1.36 | 70.80 | 6.08 | 11.65 | 437 | 533 | 30 | 1.0784 | Cambisol | Fs | Fs_pure | 50_100 |
| AEF022_17 | 6.31 | 58.86 | 0.71 | 58.15 | 4.53 | 12.84 | 542 | 409 | 49 | 0.7178 | Cambisol | Fs | Fs_pure | 30_50 |
| AEF023_17 | 5.59 | 56.94 | 0.38 | 56.55 | 4.51 | 12.53 | 500 | 486 | 15 | 1.2499 | Cambisol | Fs | Fs_pure | 50_100 |
| AEF038_17 | 6.88 | 57.30 | 1.49 | 55.81 | 4.06 | 13.75 | 502 | 387 | 111 | 1.5666 | Leptosol | Fs | Fs_pure | 15_30 |
| AEF039_17 | 5.17 | 53.14 | 0.39 | 52.75 | 3.95 | 13.36 | 432 | 530 | 38 | 1.2416 | Leptosol | Fs | Fs_pure | 15_30 |
| AEF040_17 | 5.32 | 81.85 | 0.41 | 81.43 | 6.67 | 12.22 | 610 | 347 | 44 | 0.9907 | Leptosol | Fs | Fs_pure | 50_100 |

| | | | | | | | | | | | | | | |
|---|---|---|---|---|---|---|---|---|---|---|---|---|---|---|
| AEF041_17 | 5.67 | 61.93 | 0.49 | 61.43 | 4.77 | 12.88 | 475 | 449 | 76 | 1.0057 | Leptosol | Fs | Fs_pure | 30_50 |
| AEF042_17 | 6.45 | 72.10 | 0.86 | 71.24 | 5.54 | 12.87 | 504 | 386 | 110 | 1.2601 | Leptosol | Fs | Fs_pure | 30_50 |
| AEF043_17 | 5.07 | 42.94 | 0.00 | 42.94 | 3.23 | 13.28 | 386 | 577 | 39 | 1.0001 | Leptosol | Fs | Fs_pure | 50_100 |
| AEF050_17 | 5.88 | 99.55 | 0.97 | 98.58 | 7.21 | 13.68 | 543 | 429 | 28 | 0.9005 | Leptosol | Fs | Fs_pure | 50_100 |
| HEF005_17 | 5.32 | 52.23 | 0.79 | 51.45 | 3.94 | 13.05 | 460 | 485 | 59 | 0.9556 | Luvisol | Fs | Fs_pure | 30_50 |
| HEF006_17 | 4.35 | 31.98 | 0.00 | 31.98 | 2.50 | 12.79 | 218 | 709 | 74 | 0.7467 | Luvisol | Fs | Fs_pure | 50_100 |
| HEF007_17 | 4.14 | 32.60 | 0.00 | 32.60 | 2.37 | 13.75 | 197 | 714 | 88 | 0.6948 | Luvisol | Fs | Fs_pure | 50_100 |
| HEF008_17 | 5.68 | 35.10 | 0.00 | 35.10 | 2.58 | 13.62 | 227 | 716 | 58 | 1.1562 | Luvisol | Fs | Fs_pure | 50_100 |
| HEF009_17 | 4.38 | 29.20 | 0.00 | 29.20 | 2.15 | 13.56 | 287 | 622 | 92 | 1.2574 | Luvisol | Fs | Fs_pure | 50_100 |
| HEF012_17 | 4.14 | 28.86 | 0.00 | 28.86 | 2.03 | 14.21 | 168 | 713 | 117 | 0.0000 | Luvisol | Fs | Fs_pure | 50_100 |
| HEF020_17 | 6.68 | 45.16 | 3.28 | 41.88 | 3.25 | 12.89 | 253 | 708 | 40 | 0.9926 | Luvisol | Fs | Fs_pure | 30_50 |
| HEF021_17 | 6.29 | 41.04 | 1.64 | 39.39 | 2.95 | 13.35 | 268 | 672 | 61 | 0.8144 | Luvisol | Fs | Fs_pure | 50_100 |
| HEF022_17 | 4.82 | 27.92 | 0.00 | 27.92 | 2.01 | 13.87 | 184 | 747 | 68 | 0.6386 | Luvisol | Fs | Fs_pure | 50_100 |
| HEF023_17 | 4.72 | 41.31 | 0.00 | 41.31 | 3.18 | 12.99 | 279 | 662 | 56 | 1.3847 | Luvisol | Fs | Fs_pure | 50_100 |
| HEF024_17 | 3.98 | 24.67 | 0.00 | 24.67 | 1.80 | 13.74 | 162 | 793 | 44 | 0.8871 | Luvisol | Fs | Fs_pure | 50_100 |
| HEF025_17 | 4.74 | 43.96 | 0.00 | 43.96 | 3.43 | 12.80 | 325 | 624 | 53 | 1.2635 | Luvisol | Fs | Fs_pure | 50_100 |
| HEF026_17 | 4.31 | 25.49 | 0.00 | 25.49 | 1.79 | 14.26 | 150 | 796 | 54 | 1.1968 | Luvisol | Fs | Fs_pure | 50_100 |
| HEF027_17 | 6.02 | 56.31 | 0.00 | 56.31 | 4.20 | 13.40 | 291 | 643 | 67 | 0.7979 | Luvisol | Fs | Fs_pure | 50_100 |
| HEF028_17 | 6.19 | 53.61 | 0.63 | 52.98 | 3.83 | 13.83 | 306 | 634 | 60 | 1.4438 | Stagnosol | Fs | Fs_pure | 50_100 |
| HEF029_17 | 4.12 | 33.64 | 0.00 | 33.64 | 2.48 | 13.58 | 227 | 719 | 55 | 0.5728 | Luvisol | Fs | Fs_pure | 50_100 |
| HEF030_17 | 4.06 | 37.21 | 0.00 | 37.21 | 2.94 | 12.64 | 358 | 595 | 48 | 0.7626 | Luvisol | Fs | Fs_pure | 50_100 |
| HEF031_17 | 4.13 | 42.14 | 0.00 | 42.14 | 3.22 | 13.08 | 377 | 573 | 52 | 1.0788 | Luvisol | Fs | Fs_pure | 50_100 |
| HEF032_17 | 3.93 | 49.21 | 0.00 | 49.21 | 3.60 | 13.67 | 345 | 593 | 64 | 1.1500 | Luvisol | Fs | Fs_pure | 50_100 |
| HEF033_17 | 4.80 | 39.09 | 0.00 | 39.09 | 2.50 | 15.63 | 227 | 706 | 68 | 1.1054 | Luvisol | Fs | Fs_pure | 50_100 |
| HEF037_17 | 4.41 | 36.75 | 0.00 | 36.75 | 2.67 | 13.74 | 265 | 661 | 78 | 0.0000 | Stagnosol | Fs | Fs_pure | 50_100 |
| HEF042_17 | 4.17 | 31.24 | 0.00 | 31.24 | 2.14 | 14.62 | 184 | 760 | 60 | 0.5400 | Stagnosol | Fs | Fs_pure | 50_100 |
| HEF045_17 | 7.15 | 78.62 | 9.26 | 69.36 | 5.18 | 13.39 | 88 | 854 | 60 | 1.0203 | Luvisol | Fs | Fs_pure | 15_30 |
| HEF046_17 | 4.19 | 46.55 | 0.00 | 46.55 | 3.63 | 12.81 | 387 | 570 | 45 | 1.0506 | Luvisol | Fs | Fs_pure | 30_50 |
| HEF047_17 | 4.86 | 33.95 | 0.00 | 33.95 | 2.57 | 13.22 | 323 | 632 | 46 | 0.8971 | Stagnosol | Fs | Fs_pure | 50_100 |
| HEF048_17 | 4.42 | 37.01 | 0.00 | 37.01 | 2.47 | 14.97 | 273 | 687 | 44 | 0.8371 | Stagnosol | Fs | Fs_pure | 50_100 |
| HEF049_17 | 4.07 | 30.28 | 0.00 | 30.28 | 2.02 | 14.99 | 223 | 732 | 48 | 1.2523 | Stagnosol | Fs | Fs_pure | 50_100 |
| HEF050_17 | 4.77 | 42.94 | 0.00 | 42.94 | 3.57 | 12.03 | 349 | 606 | 46 | 0.4231 | Stagnosol | Fs | Fs_pure | 50_100 |
| SEF005_17 | 3.36 | 25.40 | 0.00 | 25.40 | 1.32 | 19.23 | 19 | 17 | 964 | 0.6890 | Cambisol | Fs | Fs_pure | 50_100 |
| SEF006_17 | 3.66 | 27.93 | 0.00 | 27.93 | 1.68 | 16.59 | 49 | 77 | 874 | 1.4018 | Cambisol | Fs | Fs_pure | 50_100 |
| SEF007_17 | 3.75 | 24.22 | 0.00 | 24.22 | 1.55 | 15.65 | 16 | 124 | 860 | 0.0855 | Cambisol | Fs | Fs_pure | 50_100 |
| SEF008_17 | 3.37 | 24.19 | 0.00 | 24.19 | 1.50 | 16.15 | 47 | 198 | 755 | 0.1873 | Albeluvisol | Fs | Fs_pure | 50_100 |
| SEF009_17 | 3.49 | 16.87 | 0.00 | 16.87 | 0.91 | 18.57 | 0 | 58 | 942 | 0.6459 | Cambisol | Fs | Fs_pure | 50_100 |
| SEF035_17 | 3.64 | 19.11 | 0.00 | 19.11 | 1.04 | 18.29 | 65 | 74 | 861 | 1.0252 | Cambisol | Fs | Fs_pure | 50_100 |
| SEF036_17 | 3.29 | 30.93 | 0.00 | 30.93 | 1.77 | 17.51 | 64 | 88 | 848 | 1.2034 | Cambisol | Fs | Fs_pure | 30_50 |
| SEF039_17 | 3.73 | 19.19 | 0.00 | 19.19 | 1.18 | 16.33 | 52 | 98 | 850 | 1.0994 | Cambisol | Fs | Fs_pure | 50_100 |
| SEF040_17 | 3.77 | 21.49 | 0.00 | 21.49 | 1.26 | 17.03 | 63 | 71 | 866 | 1.3649 | Cambisol | Fs | Fs_pure | 50_100 |
| SEF041_17 | 3.78 | 21.47 | 0.00 | 21.47 | 1.22 | 17.54 | 50 | 49 | 905 | 1.1019 | Cambisol | Fs | Fs_pure | 50_100 |
| SEF042_17 | 3.75 | 24.71 | 0.00 | 24.71 | 1.50 | 16.48 | 74 | 247 | 679 | 0.7863 | Regosol | Fs | Fs_pure | 50_100 |
| SEF043_17 | 3.67 | 16.96 | 0.00 | 16.96 | 1.04 | 16.31 | 69 | 121 | 810 | 1.0317 | Cambisol | Fs | Fs_pure | 50_100 |
| SEF045_17 | 3.71 | 17.17 | 0.00 | 17.17 | 1.08 | 15.92 | 48 | 75 | 881 | 0.3413 | Albeluvisol | Fs | Fs_pure | 50_100 |
| SEF046_17 | 3.49 | 24.08 | 0.00 | 24.08 | 1.48 | 16.27 | 63 | 132 | 805 | 0.2069 | Albeluvisol | Fs | Fs_pure | 50_100 |
| SEF047_17 | 3.45 | 20.56 | 0.00 | 20.56 | 1.17 | 17.63 | 49 | 18 | 933 | 0.0156 | Cambisol | Fs | Fs_pure | 50_100 |
| SEF049_17 | 3.47 | 26.53 | 0.00 | 26.53 | 1.44 | 18.42 | 68 | 74 | 858 | 1.0371 | Cambisol | Fs | Fs_pure | 50_100 |
| SEF050_17 | 3.68 | 21.17 | 0.00 | 21.17 | 1.34 | 15.85 | 79 | 99 | 822 | 1.1073 | Cambisol | Fs | Fs_pure | 50_100 |
| SEF037_11 | 3.43 | 21.34 | 0.00 | 21.34 | 1.26 | 16.97 | 63 | 70 | 871 | 0.6602 | Cambisol | Fs | Fs_Qs_mixed | 50_100 |
| SEF044_11 | 3.52 | 17.32 | 0.00 | 17.32 | 0.94 | 18.47 | 75 | 69 | 856 | 0.7971 | Cambisol | Fs | Fs_Qs_mixed | 50_100 |
| SEF037_17 | 3.56 | 22.95 | 0.00 | 22.95 | 1.27 | 18.08 | 63 | 70 | 871 | 0.6602 | Cambisol | Fs | Fs_Qs_mixed | 50_100 |
| SEF044_17 | 3.74 | 23.31 | 0.00 | 23.31 | 1.34 | 17.34 | 75 | 69 | 856 | 0.7971 | Cambisol | Fs | Fs_Qs_mixed | 50_100 |
| AEF021_11 | 6.11 | 72.49 | 0.62 | 71.87 | 6.14 | 11.71 | 519 | 453 | 30 | 1.0136 | Cambisol | Fs | Fs_oHS_mixed | 50_100 |

| | | | | | | | | | | | | | | |
|---|---|---|---|---|---|---|---|---|---|---|---|---|---|---|
| AEF026_11 | 4.85 | 47.45 | 0.22 | 47.24 | 3.87 | 12.19 | 480 | 329 | 181 | 1.6704 | Cambisol | Fs | Fs_oHS_mixed | 15_30 |
| AEF045_11 | 5.71 | 63.39 | 0.42 | 62.96 | 5.15 | 12.22 | 220 | 574 | 206 | 1.0005 | Cambisol | Fs | Fs_oHS_mixed | 0-15 |
| HEF016_11 | 4.74 | 33.26 | 0.26 | 33.01 | 2.73 | 12.10 | 322 | 628 | 53 | 0.7930 | Luvisol | Fs | Fs_oHS_mixed | 15_30 |
| AEF021_17 | 6.31 | 71.96 | 0.81 | 71.15 | 6.08 | 11.70 | 519 | 453 | 30 | 1.0136 | Cambisol | Fs | Fs_oHS_mixed | 50_100 |
| AEF026_17 | 5.10 | 52.96 | 0.00 | 52.96 | 4.09 | 12.95 | 480 | 329 | 181 | 1.6704 | Cambisol | Fs | Fs_oHS_mixed | 15_30 |
| AEF045_17 | 5.80 | 64.12 | 0.36 | 63.76 | 5.15 | 12.37 | 220 | 574 | 206 | 1.0005 | Cambisol | Fs | Fs_oHS_mixed | 0-15 |
| HEF016_17 | 4.86 | 35.88 | 0.00 | 35.88 | 2.87 | 12.49 | 322 | 628 | 53 | 0.7930 | Luvisol | Fs | Fs_oHS_mixed | 15_30 |
| AEF024_11 | 4.88 | 57.54 | 0.30 | 57.25 | 4.39 | 13.04 | 510 | 307 | 183 | 2.5668 | Cambisol | Fs | 3_mixed | 0-15 |
| AEF024_17 | 5.29 | 58.61 | 0.30 | 58.31 | 4.55 | 12.80 | 510 | 307 | 183 | 2.5668 | Cambisol | Fs | 3_mixed | 0-15 |
| AEF029_11 | 4.49 | 51.60 | 0.24 | 51.36 | 3.69 | 13.92 | 413 | 550 | 37 | 2.0625 | Leptosol | Fs | Fs_Pa_mixed | 30_50 |
| AEF029_17 | 4.40 | 53.18 | 0.00 | 53.18 | 3.79 | 14.02 | 413 | 550 | 37 | 2.0625 | Leptosol | Fs | Fs_Pa_mixed | 30_50 |
| HEF013_11 | 6.66 | 70.65 | 1.40 | 69.25 | 4.58 | 15.13 | 510 | 444 | 48 | 2.2452 | Luvisol | Pa | Pa_oHS_mixed | 30_50 |
| HEF013_17 | 6.76 | 78.11 | 1.54 | 76.58 | 4.83 | 15.84 | 510 | 444 | 48 | 2.2452 | Luvisol | Pa | Pa_oHS_mixed | 30_50 |
| HEF001_11 | 6.78 | 49.15 | 3.58 | 45.57 | 3.63 | 12.54 | 195 | 738 | 71 | 1.8694 | Stagnosol | Pa | 3_mixed | 50_100 |
| HEF001_17 | 6.23 | 51.33 | 1.11 | 50.22 | 3.91 | 12.85 | 195 | 738 | 71 | 1.8694 | Stagnosol | Pa | 3_mixed | 50_100 |
| HEF002_11 | 5.53 | 41.63 | 0.39 | 41.24 | 2.71 | 15.23 | 308 | 600 | 92 | 1.3073 | Stagnosol | Pa | 3_mixed | 30_50 |
| HEF002_17 | 6.59 | 68.36 | 1.08 | 67.28 | 3.83 | 17.55 | 308 | 600 | 92 | 1.3073 | Stagnosol | Pa | 3_mixed | 30_50 |
| AEF001_11 | 3.43 | 56.53 | 0.24 | 56.29 | 3.67 | 15.34 | 318 | 659 | 23 | 1.8052 | Cambisol | Pa | Pa_pure | 30_50 |
| AEF002_11 | 4.34 | 53.26 | 0.23 | 53.03 | 3.81 | 13.91 | 500 | 462 | 38 | 2.2656 | Leptosol | Pa | Pa_pure | 30_50 |
| AEF003_11 | 5.66 | 69.24 | 0.72 | 68.52 | 4.83 | 14.18 | 527 | 424 | 49 | 2.3526 | Cambisol | Pa | Pa_pure | 30_50 |
| AEF010_11 | 4.63 | 57.48 | 0.24 | 57.24 | 4.24 | 13.48 | 528 | 441 | 33 | 2.1337 | Leptosol | Pa | Pa_pure | 30_50 |
| AEF011_11 | 3.49 | 54.13 | 0.24 | 53.89 | 3.57 | 15.09 | 260 | 699 | 41 | 1.6501 | Cambisol | Pa | Pa_pure | 50_100 |
| AEF012_11 | 4.29 | 60.44 | 0.29 | 60.16 | 4.57 | 13.16 | 570 | 405 | 25 | 2.3458 | Cambisol | Pa | Pa_pure | 50_100 |
| AEF013_11 | 4.76 | 63.93 | 0.27 | 63.67 | 4.29 | 14.85 | 666 | 289 | 45 | 1.7833 | Cambisol | Pa | Pa_pure | 50_100 |
| AEF014_11 | 4.59 | 42.17 | 0.27 | 41.90 | 3.27 | 12.81 | 449 | 508 | 43 | 1.7546 | Cambisol | Pa | Pa_pure | 50_100 |
| AEF031_11 | 5.81 | 70.31 | 0.59 | 69.72 | 5.71 | 12.20 | 548 | 431 | 21 | 2.2552 | Leptosol | Pa | Pa_pure | 30_50 |
| AEF032_11 | 6.58 | 105.70 | 6.68 | 99.06 | 7.64 | 12.95 | 395 | 548 | 57 | 2.3871 | Leptosol | Pa | Pa_pure | 30_50 |
| AEF033_11 | 5.33 | 87.84 | 0.52 | 87.32 | 6.06 | 14.40 | 681 | 305 | 14 | 2.3050 | Cambisol | Pa | Pa_pure | 30_50 |
| AEF034_11 | 4.93 | 70.84 | 0.32 | 70.51 | 5.15 | 13.70 | 626 | 353 | 23 | 2.2675 | Leptosol | Pa | Pa_pure | 30_50 |
| HEF003_11 | 4.98 | 43.33 | 0.33 | 43.00 | 2.72 | 15.79 | 409 | 549 | 43 | 2.1979 | Luvisol | Pa | Pa_pure | 30_50 |
| AEF001_17 | 3.34 | 72.74 | 0.34 | 72.41 | 4.47 | 16.20 | 318 | 659 | 23 | 1.8052 | Cambisol | Pa | Pa_pure | 30_50 |
| AEF002_17 | 4.84 | 53.20 | 0.32 | 52.88 | 3.82 | 13.85 | 500 | 462 | 38 | 2.2656 | Leptosol | Pa | Pa_pure | 30_50 |
| AEF003_17 | 5.63 | 64.02 | 0.56 | 63.46 | 4.67 | 13.58 | 527 | 424 | 49 | 2.3526 | Cambisol | Pa | Pa_pure | 30_50 |
| AEF010_17 | 4.63 | 47.34 | 0.33 | 47.02 | 3.77 | 12.46 | 528 | 441 | 33 | 2.1337 | Leptosol | Pa | Pa_pure | 30_50 |
| AEF011_17 | 3.42 | 60.03 | 0.30 | 59.73 | 3.92 | 15.22 | 260 | 699 | 41 | 1.6501 | Cambisol | Pa | Pa_pure | 50_100 |
| AEF012_17 | 4.52 | 58.27 | 0.37 | 57.90 | 4.49 | 12.89 | 570 | 405 | 25 | 2.3458 | Cambisol | Pa | Pa_pure | 50_100 |
| AEF013_17 | 5.16 | 59.49 | 0.31 | 59.18 | 3.99 | 14.82 | 666 | 289 | 45 | 1.7833 | Cambisol | Pa | Pa_pure | 50_100 |
| AEF014_17 | 4.84 | 46.57 | 0.00 | 46.57 | 3.49 | 13.34 | 449 | 508 | 43 | 1.7546 | Cambisol | Pa | Pa_pure | 50_100 |
| AEF031_17 | 5.59 | 61.18 | 0.44 | 60.74 | 4.97 | 12.21 | 548 | 431 | 21 | 2.2552 | Leptosol | Pa | Pa_pure | 30_50 |
| AEF032_17 | 6.93 | 114.43 | 11.03 | 103.40 | 7.72 | 13.40 | 395 | 548 | 57 | 2.3871 | Leptosol | Pa | Pa_pure | 30_50 |
| AEF033_17 | 5.79 | 92.80 | 0.87 | 91.92 | 6.06 | 15.16 | 681 | 305 | 14 | 2.3050 | Cambisol | Pa | Pa_pure | 30_50 |
| AEF034_17 | 4.90 | 72.42 | 0.38 | 72.04 | 5.15 | 14.00 | 626 | 353 | 23 | 2.2675 | Leptosol | Pa | Pa_pure | 30_50 |
| HEF003_17 | 5.07 | 59.14 | 0.47 | 58.68 | 3.56 | 16.47 | 409 | 549 | 43 | 2.1979 | Luvisol | Pa | Pa_pure | 30_50 |
| SEF004_11 | 3.27 | 24.63 | 0.00 | 24.63 | 1.19 | 20.75 | 27 | 54 | 919 | 1.5961 | Cambisol | Ps | Fs_Ps_mixed | 50_100 |
| SEF029_11 | 3.18 | 12.02 | 0.00 | 12.02 | 0.55 | 21.93 | 30 | 11 | 959 | 1.5470 | Cambisol | Ps | Fs_Ps_mixed | 50_100 |
| SEF031_11 | 3.23 | 20.53 | 0.00 | 20.53 | 1.06 | 19.37 | 51 | 61 | 888 | 1.8482 | Cambisol | Ps | Fs_Ps_mixed | 50_100 |
| SEF033_11 | 3.18 | 12.55 | 0.00 | 12.55 | 0.50 | 24.96 | 25 | 9 | 966 | 1.7008 | Cambisol | Ps | Fs_Ps_mixed | 50_100 |
| SEF004_17 | 3.46 | 24.81 | 0.00 | 24.81 | 1.24 | 19.94 | 27 | 54 | 919 | 1.5961 | Cambisol | Ps | Fs_Ps_mixed | 50_100 |
| SEF029_17 | 3.32 | 20.44 | 0.00 | 20.44 | 0.87 | 23.63 | 30 | 11 | 959 | 1.5470 | Cambisol | Ps | Fs_Ps_mixed | 50_100 |
| SEF031_17 | 3.41 | 28.22 | 0.00 | 28.22 | 1.41 | 20.03 | 51 | 61 | 888 | 1.8482 | Cambisol | Ps | Fs_Ps_mixed | 50_100 |
| SEF033_17 | 3.35 | 11.64 | 0.00 | 11.64 | 0.48 | 24.18 | 25 | 9 | 966 | 1.7008 | Cambisol | Ps | Fs_Ps_mixed | 50_100 |
| SEF001_11 | 3.50 | 18.60 | 0.00 | 18.60 | 0.80 | 23.20 | 37 | 81 | 882 | 1.9003 | Cambisol | Ps | Ps_pure | 15_30 |
| SEF002_11 | 3.40 | 20.56 | 0.00 | 20.56 | 1.20 | 17.07 | 32 | 98 | 870 | 1.4052 | Cambisol | Ps | Ps_pure | 30_50 |
| SEF003_11 | 3.34 | 20.54 | 0.00 | 20.54 | 1.03 | 19.84 | 24 | 50 | 926 | 1.8981 | Cambisol | Ps | Ps_pure | 30_50 |

| | | | | | | | | | | | | | | |
|---|---|---|---|---|---|---|---|---|---|---|---|---|---|---|
| SEF010_11 | 3.58 | 18.31 | 0.00 | 18.31 | 1.02 | 17.94 | 36 | 53 | 911 | 1.1101 | Cambisol | Ps | Ps_pure | 15_30 |
| SEF011_11 | 3.29 | 31.32 | 0.00 | 31.32 | 1.31 | 23.83 | 31 | 85 | 884 | 2.1194 | Cambisol | Ps | Ps_pure | 15_30 |
| SEF012_11 | 3.38 | 20.76 | 0.00 | 20.76 | 0.79 | 26.30 | 18 | 106 | 876 | 2.2966 | Cambisol | Ps | Ps_pure | 15_30 |
| SEF013_11 | 3.16 | 26.57 | 0.00 | 26.57 | 1.29 | 20.53 | 28 | 51 | 921 | 1.1203 | Podzol | Ps | Ps_pure | 30_50 |
| SEF014_11 | 3.18 | 25.28 | 0.00 | 25.28 | 1.43 | 17.63 | 8 | 17 | 975 | 1.7147 | Cambisol | Ps | Ps_pure | 30_50 |
| SEF015_11 | 3.46 | 10.68 | 0.00 | 10.68 | 0.55 | 19.57 | 31 | 46 | 923 | 1.7954 | Cambisol | Ps | Ps_pure | 30_50 |
| SEF016_11 | 3.41 | 19.44 | 0.00 | 19.44 | 1.06 | 18.28 | 30 | 108 | 862 | 1.1101 | Cambisol | Ps | Ps_pure | 50_100 |
| SEF017_11 | 3.08 | 21.75 | 0.00 | 21.75 | 0.92 | 23.57 | 24 | 49 | 927 | 1.4928 | Cambisol | Ps | Ps_pure | 50_100 |
| SEF018_11 | 3.42 | 31.35 | 0.00 | 31.35 | 1.43 | 21.97 | 29 | 34 | 941 | 1.7025 | Cambisol | Ps | Ps_pure | 30_50 |
| SEF019_11 | 3.51 | 6.34 | 0.00 | 6.34 | 0.31 | 20.17 | 37 | 44 | 919 | 2.1162 | Cambisol | Ps | Ps_pure | 30_50 |
| SEF020_11 | 3.59 | 16.34 | 0.00 | 16.34 | 0.69 | 23.59 | 42 | 70 | 892 | 1.8772 | Regosol | Ps | Ps_pure | 30_50 |
| SEF021_11 | 3.15 | 25.43 | 0.00 | 25.43 | 1.37 | 18.60 | 55 | 69 | 876 | 1.9226 | Cambisol | Ps | Ps_pure | 50_100 |
| SEF001_17 | 3.64 | 23.28 | 0.00 | 23.28 | 1.05 | 22.18 | 37 | 81 | 882 | 1.9003 | Cambisol | Ps | Ps_pure | 15_30 |
| SEF002_17 | 3.52 | 25.01 | 0.00 | 25.01 | 1.47 | 16.98 | 32 | 98 | 870 | 1.4052 | Cambisol | Ps | Ps_pure | 30_50 |
| SEF003_17 | 3.44 | 21.41 | 0.00 | 21.41 | 1.03 | 20.76 | 24 | 50 | 926 | 1.8981 | Cambisol | Ps | Ps_pure | 30_50 |
| SEF010_17 | 3.72 | 23.46 | 0.00 | 23.46 | 1.23 | 19.07 | 36 | 53 | 911 | 1.1101 | Cambisol | Ps | Ps_pure | 15_30 |
| SEF011_17 | 3.67 | 16.08 | 0.00 | 16.08 | 0.77 | 20.99 | 31 | 85 | 884 | 2.1194 | Cambisol | Ps | Ps_pure | 15_30 |
| SEF012_17 | 3.50 | 22.62 | 0.00 | 22.62 | 0.90 | 25.24 | 18 | 106 | 876 | 2.2966 | Cambisol | Ps | Ps_pure | 15_30 |
| SEF013_17 | 3.25 | 26.44 | 0.00 | 26.44 | 1.25 | 21.15 | 28 | 51 | 921 | 1.1203 | Podzol | Ps | Ps_pure | 30_50 |
| SEF014_17 | 3.38 | 24.38 | 0.00 | 24.38 | 1.41 | 17.30 | 8 | 17 | 975 | 1.7147 | Cambisol | Ps | Ps_pure | 30_50 |
| SEF015_17 | 3.68 | 10.82 | 0.00 | 10.82 | 0.63 | 17.14 | 31 | 46 | 923 | 1.7954 | Cambisol | Ps | Ps_pure | 30_50 |
| SEF016_17 | 3.60 | 14.97 | 0.00 | 14.97 | 0.84 | 17.75 | 30 | 108 | 862 | 1.1101 | Cambisol | Ps | Ps_pure | 50_100 |
| SEF017_17 | 3.34 | 20.39 | 0.00 | 20.39 | 0.90 | 22.60 | 24 | 49 | 927 | 1.4928 | Cambisol | Ps | Ps_pure | 50_100 |
| SEF018_17 | 3.31 | 27.44 | 0.00 | 27.44 | 1.28 | 21.38 | 29 | 34 | 941 | 1.7025 | Cambisol | Ps | Ps_pure | 30_50 |
| SEF019_17 | 3.64 | 15.63 | 0.00 | 15.63 | 0.75 | 20.94 | 37 | 44 | 919 | 2.1162 | Cambisol | Ps | Ps_pure | 30_50 |
| SEF020_17 | 3.64 | 20.65 | 0.00 | 20.65 | 0.93 | 22.21 | 42 | 70 | 892 | 1.8772 | Regosol | Ps | Ps_pure | 30_50 |
| SEF021_17 | 3.29 | 23.49 | 0.00 | 23.49 | 1.23 | 19.06 | 55 | 69 | 876 | 1.9226 | Cambisol | Ps | Ps_pure | 50_100 |
| SEF022_11 | 3.37 | 23.37 | 0.00 | 23.37 | 1.31 | 17.86 | 34 | 71 | 895 | 0.7747 | Cambisol | Qs | Fs_Qs_mixed | 30_50 |
| SEF023_11 | 3.24 | 19.62 | 0.00 | 19.62 | 1.30 | 15.10 | 51 | 76 | 873 | 0.9618 | Cambisol | Qs | Fs_Qs_mixed | 50_100 |
| SEF027_11 | 3.30 | 18.20 | 0.00 | 18.20 | 1.14 | 15.93 | 51 | 139 | 810 | 1.1539 | Cambisol | Qs | Fs_Qs_mixed | 50_100 |
| SEF038_11 | 3.18 | 21.87 | 0.00 | 21.87 | 1.37 | 15.96 | 57 | 103 | 840 | 0.7720 | Cambisol | Fs | Fs_Qs_mixed | 50_100 |
| SEF022_17 | 3.56 | 24.94 | 0.00 | 24.94 | 1.42 | 17.54 | 34 | 71 | 895 | 0.7747 | Cambisol | Qs | Fs_Qs_mixed | 30_50 |
| SEF023_17 | 3.48 | 18.42 | 0.00 | 18.42 | 1.28 | 14.44 | 51 | 76 | 873 | 0.9618 | Cambisol | Qs | Fs_Qs_mixed | 50_100 |
| SEF027_17 | 3.43 | 17.80 | 0.00 | 17.80 | 1.11 | 15.97 | 51 | 139 | 810 | 1.1539 | Cambisol | Qs | Fs_Qs_mixed | 50_100 |
| SEF038_17 | 3.37 | 22.81 | 0.00 | 22.81 | 1.40 | 16.24 | 57 | 103 | 840 | 0.7720 | Cambisol | Fs | Fs_Qs_mixed | 50_100 |
| SEF025_11 | 3.44 | 20.69 | 0.00 | 20.69 | 1.32 | 15.72 | 63 | 77 | 860 | 0.8530 | Cambisol | Qs | Qs_pure | 30_50 |
| SEF026_11 | 3.52 | 17.62 | 0.00 | 17.62 | 1.16 | 15.18 | 52 | 66 | 886 | 0.3760 | Cambisol | Qs | Qs_pure | 50_100 |
| SEF028_11 | 3.36 | 24.27 | 0.00 | 24.27 | 1.64 | 14.76 | 51 | 141 | 808 | 1.0689 | Cambisol | Qs | Qs_pure | 50_100 |
| SEF025_17 | 3.72 | 21.85 | 0.00 | 21.85 | 1.37 | 15.93 | 63 | 77 | 860 | 0.8530 | Cambisol | Qs | Qs_pure | 30_50 |
| SEF026_17 | 3.67 | 20.00 | 0.00 | 20.00 | 1.33 | 15.01 | 52 | 66 | 886 | 0.3760 | Cambisol | Qs | Qs_pure | 50_100 |
| SEF028_17 | 3.52 | 18.04 | 0.00 | 18.04 | 1.25 | 14.38 | 51 | 141 | 808 | 1.0689 | Cambisol | Qs | Qs_pure | 50_100 |

1: in sites SEF010 and SEF016 (2011 and 2017) , missing values were replaced by the median (1.1101). The index is is the sum of the estimation of three components: i) the proportion of harvested tree volume; ii) the proportion of invasive tree species; and iii) the proportion of dead wood showing signs of saw cuts.

2: We assembled some of the 23 categories. For two-species stands, categories sp1-sp2 and sp2-sp1 were joined. One category was made for stands with three species, thus reducing the categories to ten.

3: mean of the trunk diameter at breast height of the 100 largest trees of the main tree species
