## Supplementary material for "Distinct responses of oomycete plant parasites according to their lifestyle in a landscape-scale metabarcoding survey": Table S6

**Table S6**. Most parsimonious models (dbRDA), with their respective $R^2$ adjusted and F values, and the F values of the factors selected in each model. Significance values shown as symbol (see footnote).

### OOMYCOTA

| **Grassland** | **$R^2$ (%)** | **Anova F value** | **soil type** | **pH** | **sand** | **organic C** | **LUI** | **mowing** | **C/N ratio** |
|---|---|---|---|---|---|---|---|---|---|
| **All OTUs** | 2.6 | 1.9 *** | 1.3 * | | | | | | |
| **hemibiotrophs** | 3.3 | 2.1 *** | 1.5 * | | | | | | 2.3 * |
| **obligate biotrophs** | 3.3 | 2.3 *** | 2.3 ** | | | | 2.6 * | | |
| **saprotrophs** | 2.7 | 2.0 *** | 2.0 ** | | | | | | 2.7 ** |

| **Grassland by region** | **$R^2$ (%)** | **Anova F value** | **soil type** | **pH** | **sand** | **organic C** | **LUI** | **mowing** | **C/N ratio** |
|---|---|---|---|---|---|---|---|---|---|
| Alb | 2.6 | 1.7 * | | | | | 2.9 * | 3.4 ** | |
| Hainich | NS | NS | | | | | | | |
| Schorfheide | 5.9 | 2.0 *** | 2.1 ** | | | 2.9 * | | | 5.1 ** |

| **Forest** | **$R^2$ (%)** | **Anova F value** | **soil type** | **main tree species** | **pH** | **sand** | **organic C** | **intensity manag.** | **develop mental stage** | **C/N ratio** |
|---|---|---|---|---|---|---|---|---|---|---|
| **All OTUs** | 25.3 | 9.4 *** | 1.9 ** | 4.9 ** | 3.8 ** | 10.0** | | 2.7 ** | | |
| **hemibiotrophs** | 15.7 | 5.6 *** | 2.1 ** | 3.0 ** | 8.8 ** | 7.4 ** | | 4.3 ** | | |
| **obligate biotrophs** | 14.8 | 5.3 *** | 1.4 * | 2.7 ** | | 5.6 ** | | | | 2.0 * |
| **saprotrophs** | 38.5 | 12.0 *** | 3.3 ** | 4.0 ** | | 17.9 ** | 3.1 * | 2.7 * | 2.1 ** | 2.2 * |

| **Forest by region** | **$R^2$ (%)** | **Anova F value** | **soil type** | **main tree species** | **pH** | **sand** | **organic C** | **intensity manag.** | **develop mental stage** | **C/N ratio** |
|---|---|---|---|---|---|---|---|---|---|---|
| Alb | 11.3 | 4.2 *** | 2.9 * | 1.9 * | 4.4 ** | | | 4.4 ** | | |
| Hainich | 10.5 | 4.9 *** | | | 2.6 * | | 3.2 ** | 3.6 ** | | |
| Schorfheide | 16.7 | 2.7 *** | 1.7 * | | 4.2 ** | 3.3 ** | | 2.1 * | 2.3 ** | 2.2 * |

*p values signification codes: *** ≤ 0.001; ** ≤ 0.01; * ≤ 0.05*
